# Age-related dynamics of DNA co-methylation modules in humans

**DOI:** 10.64898/2026.09.13.751212

**Authors:** Evans K Cheruiyot, Alesha A Hatton, Daniel L McCartney, Riccardo E Marioni, Allan F McRae

**Affiliations:** Institute of Molecular Bioscience, The University of Queensland, Brisbane, QLD, Australia; Institute of Genetics and Cancer, University of Edinburgh, Edinburgh, UK

## Abstract

Age-related changes in the mean or variance of DNA methylation (DNAm) at individual CpG sites are well established. However, CpGs that share biological functions often exhibit coordinated regulation. Here, we analysed whole-blood DNAm data from over 7,500 individuals and identified modules of co-methylated CpGs that either maintain consistent correlations with age (age-stable modules) or undergo age-related remodelling (age-variable modules). We show that co-methylation modules are largely preserved during healthy aging, with only a small subset (∼15%) showing substantial age-related reorganisation.

Age-stable modules are enriched for CpG loci within promoters, transcription start sites, and CpG islands—whereas age-variable modules harbour loci that are depleted in these genomic domains and enriched for repressive chromatin marks. Age-stable co-methylation modules are also associated with long-range genetic regulation (trans-meQTLs), promoter-associated chromatin states, and higher-order chromatin interactions. While both age-stable and age-variable modules are enriched for developmental pathways, immune-related processes are distinctly associated with age-stable modules.

Overall, our findings suggest that epigenetic aging is characterised by a spectrum of co-methylation network trajectories, ranging from stability to disruption, shaped by underlying genomic regulatory architecture.

## Introduction

Aging is accompanied by widespread molecular alterations, among which changes in DNA methylation (DNAm) have been extensively described e.g., ^1,2^. Age-associated shifts in DNAm have been linked to inter-individual differences in physiological decline, disease susceptibility and mortality risk ^2,3^.

Numerous studies have characterised age-associated DNAm changes at individual CpG sites. Age-related differentially methylated positions (aDMPs) exhibit shifts in mean DNAm levels with age and form the basis of epigenetic clocks used for prediction of chronological ^4,5^ and biological age ^6^. These aDMPs show either increasing or decreasing trends in mean DNAm levels with age. Beyond mean-level shifts, age-related variably methylated positions (aVMPs) capture CpG sites where inter-individual variability in DNAm increases with age ^7^. Together, these single-CpG approaches have provided important insights into the biology of aging.

CpGs in close proximity are often correlated (co-methylated), largely driven by shared genetic regulation, chromatin architecture, or coordinated transcriptional activity ^8-10^. The predictive value of coordinated methylation features has been demonstrated by some studies e.g., ^11^ showing that combining adjacent CpG sites can improve age prediction accuracy relative to single-CpG approaches. Recent studies have linked co-methylation modules to biological traits such as longevity ^12^ and neurodegenerative diseases ^13-15^.

Previous studies have shown that various co-methylation structures are conserved across tissues and developmental stages e.g., ^16,17^. Short-term longitudinal studies have reported that co-methylation patterns remain relatively stable from birth to adolescence ^18,19^. However, it is still unclear whether age-related DNAm changes occur as coordinated reorganisation of co-methylation modules or as largely independent alterations at individual CpG sites, and which modules (if any) retain stable organisation across adulthood and late life.

Here, we investigated genome-wide co-methylation patterns across the adult lifespan to identify age-stable and age-variable co-methylation modules. We further examined functional and regulatory properties underlying these patterns.

## Results

### Age-related changes in pairwise co-methylation relationships

To investigate how co-methylation patterns change with age, we analysed a large DNA methylation (DNAm) dataset of 7,553 unrelated individuals aged 17 to 99 years in the Generation Scotland cohort (Fig. 1) ^20,21^. We divided participants into eight similar sized age groups (952–986 individuals per group; Fig. 1 and Fig. S1) to ensure comparable statistical precision in the estimation of co-methylation correlations across age strata.

**Figure 1.**
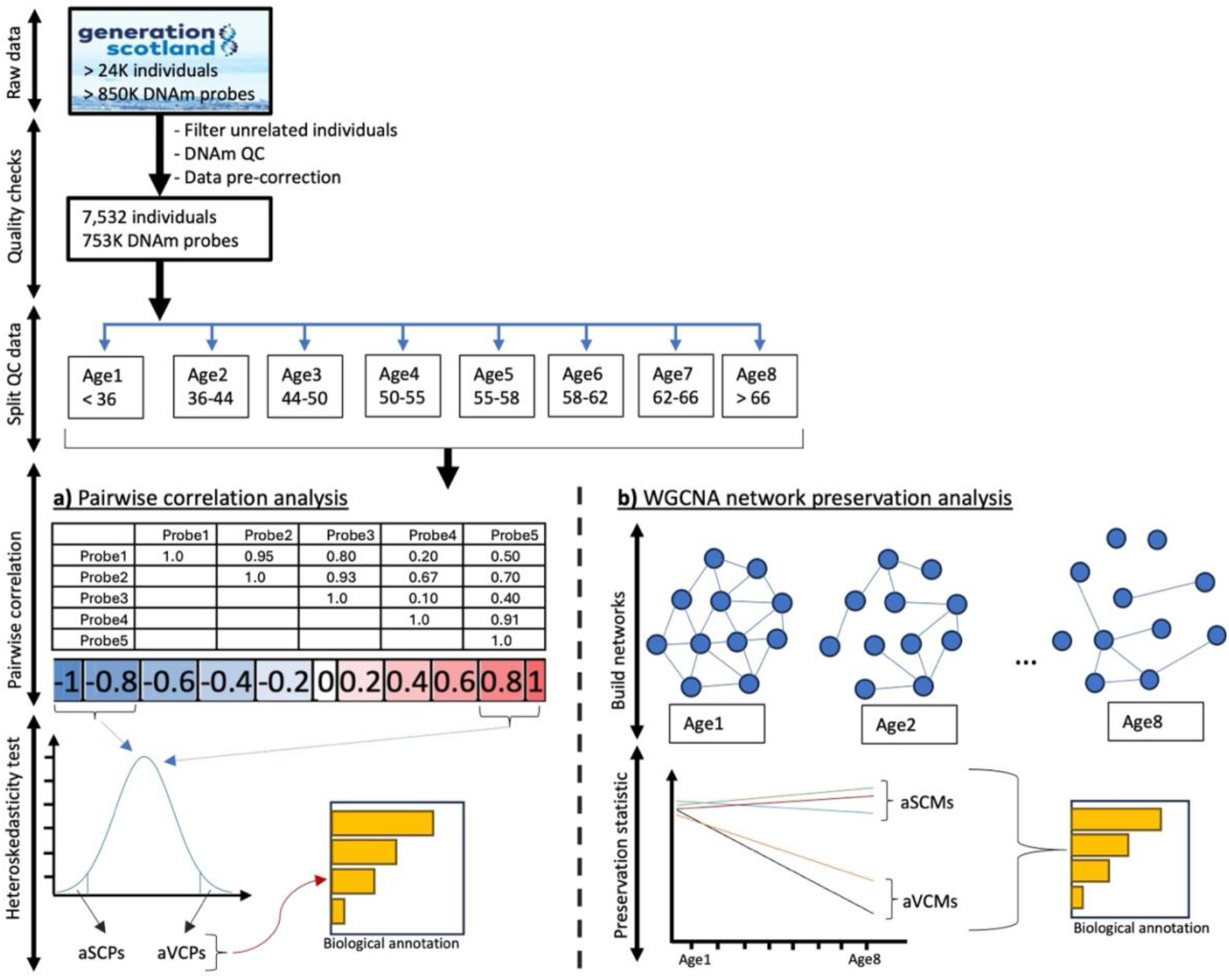
Study overview. Raw DNA methylation (DNAm) data from the Generation Scotland cohort were pre-processed, resulting in the selection of 7,532 unrelated individuals and were divided into eight age groups of comparable sample size. **(a)** Pairwise correlation analyses were conducted within each autosomal chromosome to assess age-related changes in co-methylation between CpG pairs. The Breusch-Pagan test of heteroskedasticity was then performed to identify age-variable co-methylated pairs (aVCPs). **(b)** Weighted Gene Co-expression Network Analysis (WGCNA) ^22^ was used to identify modules of co-methylated (correlated) CpGs and network preservation analysis ^23^ was performed to identify age-stable co-methylation modules (aSCMs) and age-variable co-methylation modules (aVCMs). Finally, the selected CpG sets and modules were characterised to understand their regulatory architecture and functional relevance.

We first assessed pairwise correlations (co-methylation) between CpG sites within each chromosome across age groups 1 (youngest) to 8 (oldest). Using a threshold of |r| > 0.80 in at least one age group, we identified 148,332 unique co-methylated probe pairs (Fig. S2) across the genome. A subset of these pairs (9%; N = 14,928; Fig. 2 and Fig. S2) showed significant age-related variability, based on a heteroskedasticity test (Bonferroni-adjusted p < 0.05; Fig. S2; Table S2), hereafter referred to as age-variable co-methylated probe pairs (aVCPs; see Supplementary Table S1 for a glossary of abbreviations). In contrast, the remaining co-methylated relationships remained largely preserved across age groups, here termed age-stable co-methylated probe pairs (aSCPs). aVCPs were enriched among CpG sites (OR = 3.0; p = 6.4 ×10^−8^) and genes (OR = 4.3; p = 4.18 ×10^−72^) that have been previously ^7^ associated with increased inter-individual methylation variability during aging. In contrast, aSCPs features showed no such enrichment (Table S4). Functional enrichment analysis further showed that genes associated with aVCPs were enriched for developmental processes, whereas no significant biological themes were identified for aSCPs (Fig. S4d; Table S15). Detailed genomic characteristics of these age-related pairwise co-methylation features are provided in the Supplementary Results (Supplementary Figs. S3–S8).

**Figure 1.**
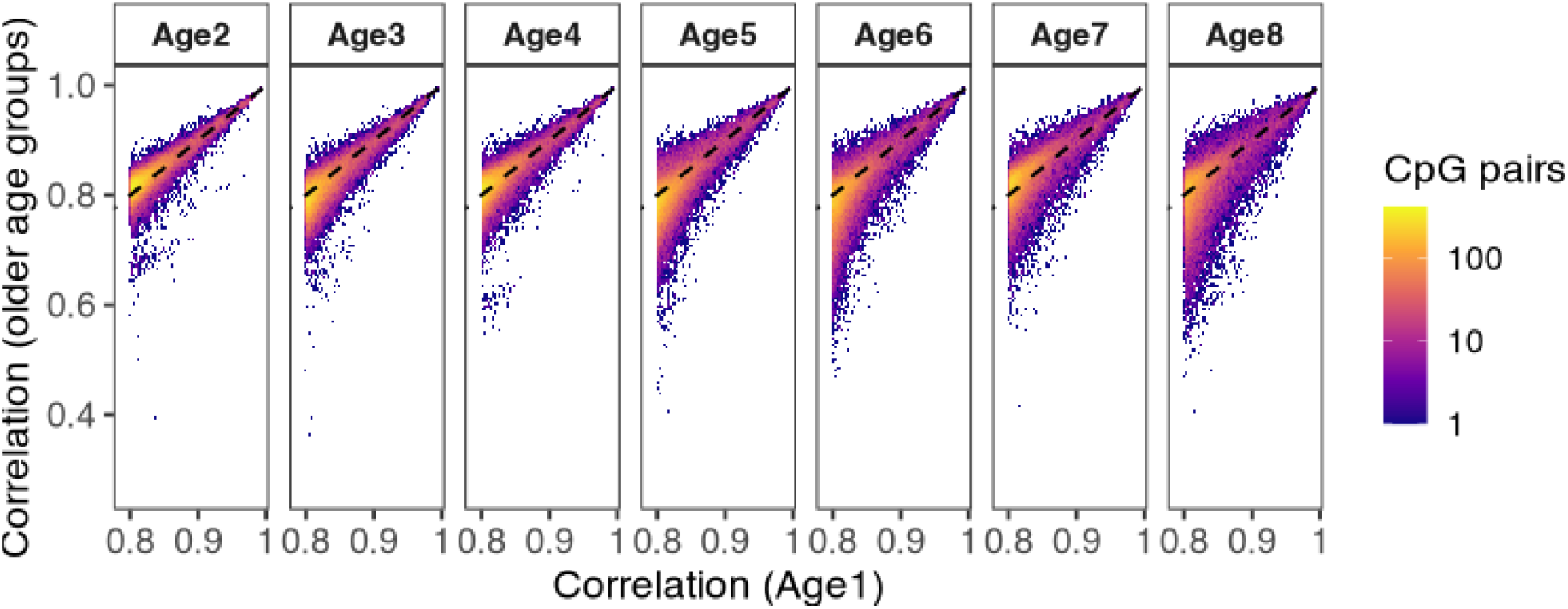
Density heatmaps of co-methylated probe pairs across different age-groups. Correlations values of positively (r > 0.80) co-methylated probe pairs identified in the youngest age group (Age1) and tracked across older age groups. Each panel compares correlation values in Age1 with those observed in subsequent age groups. Colour intensity represents the density of CpG probe pairs within each bin.

### Discovery of age-stable and age-variable co-methylation modules

To examine DNAm structure beyond pairwise correlations, we next evaluated co-methylation relationships at the network level (i.e., groups of multiple correlated CpG sites). Using the Weighted Correlation Network Analysis (WGCNA) framework ^22^, we constructed consensus co-methylation modules based on the genome-wide DNAm data from the youngest age group (reference set: age < 36 years) and quantified their preservation ^23^ across older age groups. To capture distinct aspects of age-related module dynamics, we calculated four module preservation statistics comprising one variance-based metric (proportion of variance explained) and connectivity-based metrics (cor.kIM, cor.kME and cor.cor). These three connectivity-based metrics were highly correlated (pairwise r > 0.8; Fig. S8a). As such, we selected cor.kME, which quantifies the preservation of intramodular connectivity (module hub structure), together with proportion of variance explained, which quantifies how well the module eigengene captures variation among module CpGs for downstream analyses.

Of the 69 modules identified from WGCNA (Fig. 3a; Table S4), we excluded those with a large number of CpGs (>1,000) because these modules likely capture broad background correlation structure instead of discrete co-methylation modules. Consequently, a total of 53 modules remained for downstream analysis. We applied a combination of predefined selection criteria (see Methods; Fig. S9) to distinguish age-stable from age-variable modules. Using these criteria, 11 modules showed consistent and substantial (>5%) declines in preservation between the youngest and oldest age groups (Fig. 3b and Fig. S8-9) and were classified as age-variable co-methylation modules (aVCMs). In contrast, 19 modules showed relatively stable preservation across age groups and were classified as age-stable co-methylation modules (aSCMs). Notably, our initial screening using generalized additive models (GAMs; see Methods; Fig. S9) indicated that the majority of modules (∼70%) did not show significant (p > 0.05) age-associated preservation trends. Overall, these observations suggest that co-methylation modules are largely preserved during healthy aging. The selected aSCMs comprised 675 CpGs linked to 248 genes across 19 modules (median 25 CpGs per module, range 20–74; median 6 genes per module, range 2–96), whereas aVCMs comprised 546 CpGs linked to 199 genes across 11 modules (median 46 CpGs per module, range 21–92; median 4 genes per module, range 2–134). Overlap of CpGs across preservation metrics and module sets is shown in Fig. S8.

**Figure 1.**
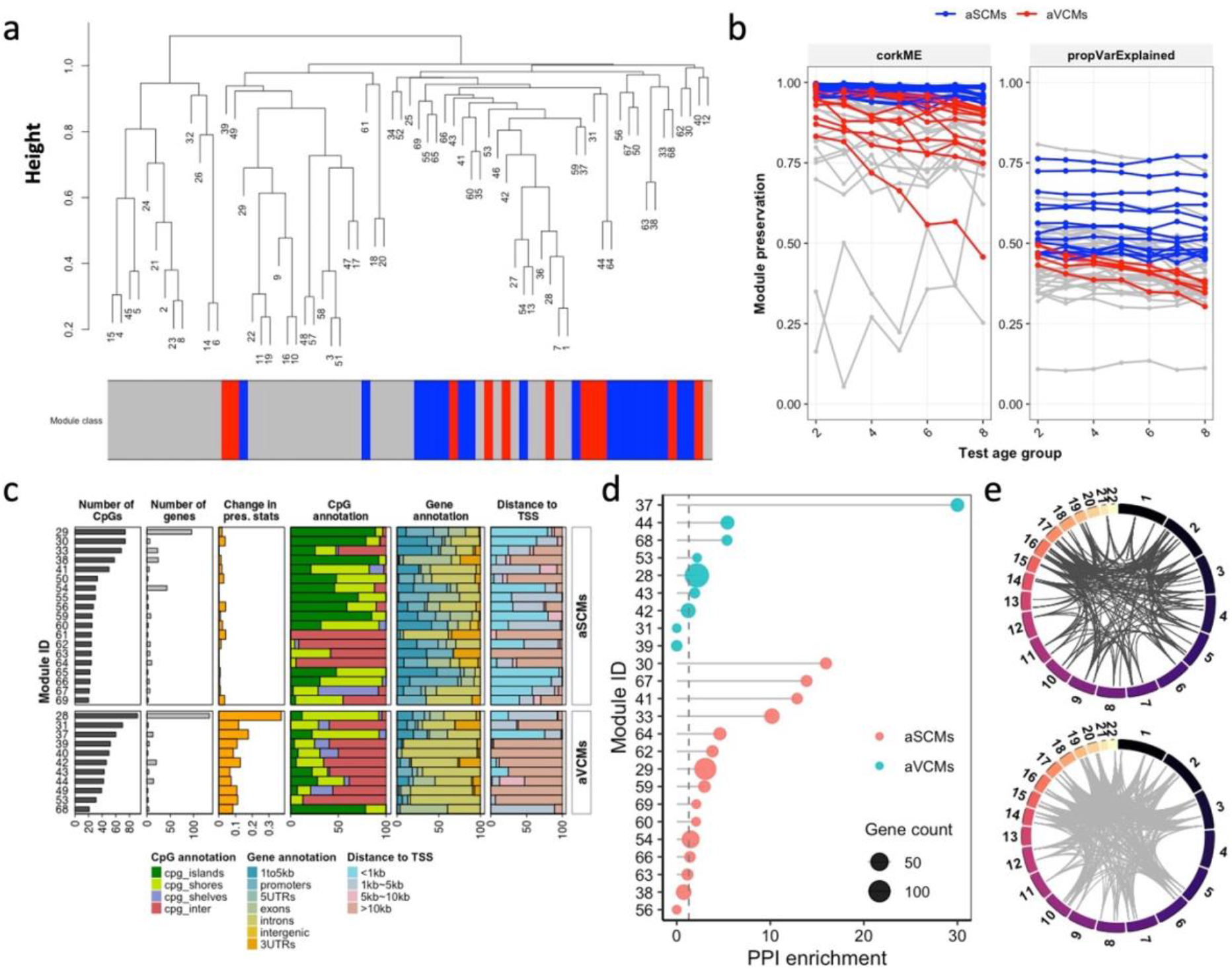
Discovery of age-stable and age-variable co-methylation modules (aSCMs and aVCMs) based on weighted gene co-methylation network analysis (WGCNA). **(a)** Co-methylation module eigengene modules constructed from genome-wide CpGs in the youngest age group (<36 years). The numbers represent module identifiers (IDs).The bar below represents aSCMs (blue) and aVCMs (red) module sets. **(b)** Preservation profiles of the selected modules classified as age-stable (aSCMs, blue) and age-variable (aVCMs, red), based on changes in either module connectivity (cor.kME) and proportion of variance explained (propVarExplained) across age groups. Grey lines represent modules not assigned to either class. **(c)** Genomic annotation profiles for aSCMs and aVCMs, depicting their distribution across various regulatory regions. **(d)** Protein–protein interaction (PPI) network enrichment (−log_10_P-value) per module for aSCMs and aVCMs (restricted to modules with >=3 genes). The vertical dashed line indicates the significance threshold (P = 0.05). **(e)** PPI modules for representative modules with the largest number of annotated genes in aSCMs (Module 28; N = 96; upper) and aVCMs (Module 29; N = 134; lower). Abbreviations: age-stable co-methylation modules (aSCMs), age-variable co-methylation modules (aVCMs).

We evaluated protein–protein interaction (PPI) connectivity enrichment for genes linked to each module set using the STRING database ^24^. Restricting analyses to modules with at least three genes, we observed that over half of aSCMs and aVCMs features show nominal significant (p < 0.05) PPI enrichment (Fig. 3d).

CpGs within aSCMs are primarily located near transcription start sites (TSSs), promoter regions, and CpG islands, whereas CpGs within the aVCMs show comparatively weaker associations within these regulatory domains (Fig. 3c and Fig. S10; Table S5–S7).

### Genetic and epigenomic architecture of co-methylation modules

We investigated whether aSCMs and aVCMs features differ in the extent to which they are influenced by genetic and environmental factors. First, to assess genetic regulation architecture, we overlapped CpGs within each module set with methylation quantitative trait loci (meQTLs) from the GoDMC project ^9^. Because the GoDMC resource was generated using the Illumina HumanMethylation450 array, analyses were restricted to CpGs represented on the 450K array. This excluded 125 (18.5%) and 214 (39.2%) CpGs from aSCMs and aVCMs, respectively. We found that both age-stable and age-variable co-methylation modules were enriched for trans- and cis-meQTLs (p < 0.001; Fig. 4a and Table S9).

**Figure 1.**
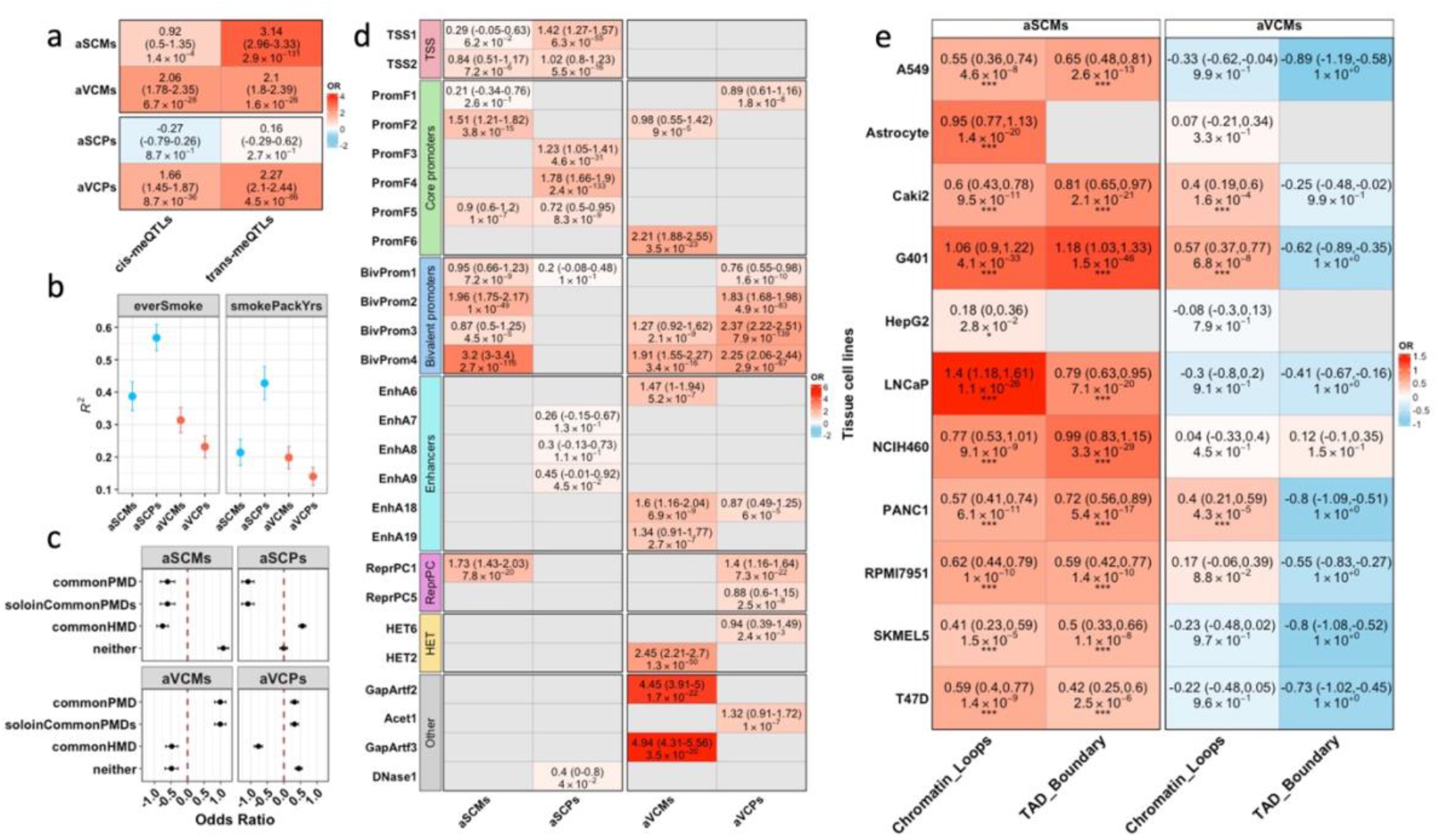
Enrichment of age-related co-methylation features across genetic, environmental, and chromatin domains. **(a)** Enrichment of stable and changing CpG sites for methylation quantitative trait loci (meQTLs) based on data from the Genetics of DNA Methylation Consortium (GoDMC ^9^). The point estimate is the log odds where > 0 represent over-enrichment and the values in brackets indicate lower and upper confidence intervals for the log odds ratio. **(b)** Proportion of variance (R^2^) in smoking status explained by age-stable and age-variable CpG sets, estimated using OSCA ^25^. The horizontal lines show the confidence intervals. **(c)** Enrichment of aSCMs and aVCMs from WGCNA in partially methylated domains (PMDs), solo WCGW CpGs, highly methylated domains (HMDs), or none of these domains, based on annotations from ^26^. The odds ratios (and CI) are represented in log scale. **(d)** Chromatin state enrichment. Grey cells denote absence of overlapping CpGs between a given CpG set and chromatin state. The cells are coloured based on odds ratio (in natural log scale). The odds ratios are represented in log scale. **(e)** Enrichment of age-stable (aSCMs) and age-variable (aVCMs) modules within higher-order chromatin loop anchors and topological associating domain (TAD) boundaries. The bars are coloured based on log(OR). The value in brackets represent the confidence intervals for the OR (in log scale) and the asterisks are the significance strength (* = p < 0.05; ** p < 0.01; *** = p < 0.001). Grey cells denote absence of overlap. Abbreviations: age-stable co-methylation modules (aSCMs), age-variable co-methylation modules (aVCMs), age-stable co-methylated probe pairs (aSCPs), age-variable co-methylated probe pairs (aVCPs).

Notably, CpGs within age-stable modules showed strong enrichment for trans-meQTLs (OR = 3.14; p = 2.9 × 10 ^−131^ ), with comparatively modest enrichment for cis-meQTLs (OR = 0.92; p = 1.0 × 10^−4^). In contrast, aVCMs showed a balanced enrichment for both cis- and trans-meQTLs (Fig. 4a), indicating that both proximal (cis) and distal genetic influences contribute to the architecture of these modules. The relative contribution of cis- and trans-meQTLs-associated CpGs varied across age-related modules (Table S10).

aSCMs explained a greater proportion of the variance (R^2^) in smoking status (ever smoker; 39%, 95% CI 34–43%) than aVCMs (31%, 95% CI 27–35%) (Fig. 4b). A similar pattern was observed for pairwise co-methylation features, with aSCPs explaining substantially more variance (57%, 95% CI 52–60%) than aVCPs (23%, 95% CI 20–26%). One possible explanation for these findings is that sustained environmental exposures, such as smoking, reinforce coordinated methylation responses across susceptible loci.

To further investigate the regulatory context of age-related co-methylation features, we tested their enrichment across several epigenomic annotations. First, we overlapped CpGs within each age-related module set with the human chromatin state annotations reported by Vu and Ernst ^25^, derived from >1,000 datasets across >100 cell types. Notably, we found that CpGs linked to the aSCMs showed enrichment for active TSS (TSS2; p = 7.2 × 10^−6^) and core promoter states (PromF2; p = 3.8 × 10^−15^) (Fig. 4d and Table S6). In contrast, aVCMs lacked enrichment across most active chromatin states but were overrepresented in constitutive heterochromatin (HET2; p = 3.5 × 10 ^−50^ ). Both aSCMs and aVCMs showed enrichment (p < 0.001) in bivalent promoter regions. In addition, aSCMs showed significant overlap with binding sites for the Polycomb regulator *CBX8* (p = 2.4 × 10?□□□; Fig. S13 and Table S7), while several additional TFBS enrichments were observed among pairwise aSCPs and aVCPs co-methylation features (Fig. S13). Moreover, aVCMs are enriched (p < 0.001) within late-replicating genomic domains (Fig. 4c and Table S8).

Age-related chromatin accessibility changes were enriched for pairwise co-methylation features (aSCPs and aVCPs) within differentially accessible regions across multiple tissues, whereas network-level modules showed weaker and tissue-specific enrichment (Supplementary Results; Fig. S14 and Table S11).

Given the enrichment of aSCMs for trans-meQTLs, we next tested whether CpGs within this module set localise within higher-order chromatin structures. Age-related co-methylation sets were overlapped with chromatin loop annotation resource from Wang, et al. ^26^ that were assayed based on Hi-C datasets from 14 selected cell lines generated by the ENCODE Consortium ^27^. The aSCMs showed the strongest and most consistent enrichment (p < 0.001) for chromatin loop regions across most cell lines (Fig. 4e; Table S12). In contrast, the age-variable module set was generally depleted (log(OR) < 1.0) for chromatin loop domains, except in three cancer-derived lines. Interestingly, we also found consistent findings when examining enrichment of age-related modules for topologically associating domains (TAD) boundaries (Fig. 4e and Supplementary Results; Table S13).

To further characterise the regulatory environments for age-related co-methylation modules, CpG linked to each module set were intersected with seven core histone modifications from the Roadmap Epigenomics Project that were profiled across 16 tissue types ^28^. We observed that both aSCMs and aVCMs were enriched for repressive histone marks, but with distinct patterns. Notably, CpGs within aSCMs showed consistent enrichment (p < 0.001) for H3K27me3 across most tissues (Fig. 5; Table S14)—a Polycomb-associated mark of facultative heterochromatin implicated in regulated gene silencing. Also, aSCMs showed enrichment for active promoter- and enhancer-associated marks, including H3K4me1 and H3K4me3. In contrast, aVCMs were enriched (p < 0.001) for H3K9me3 (a hallmark of constitutive heterochromatin linked to stable structural gene silencing) and showed no (OR < 1.0) enrichment for active histone modifications across most tissues.

**Figure 1.**
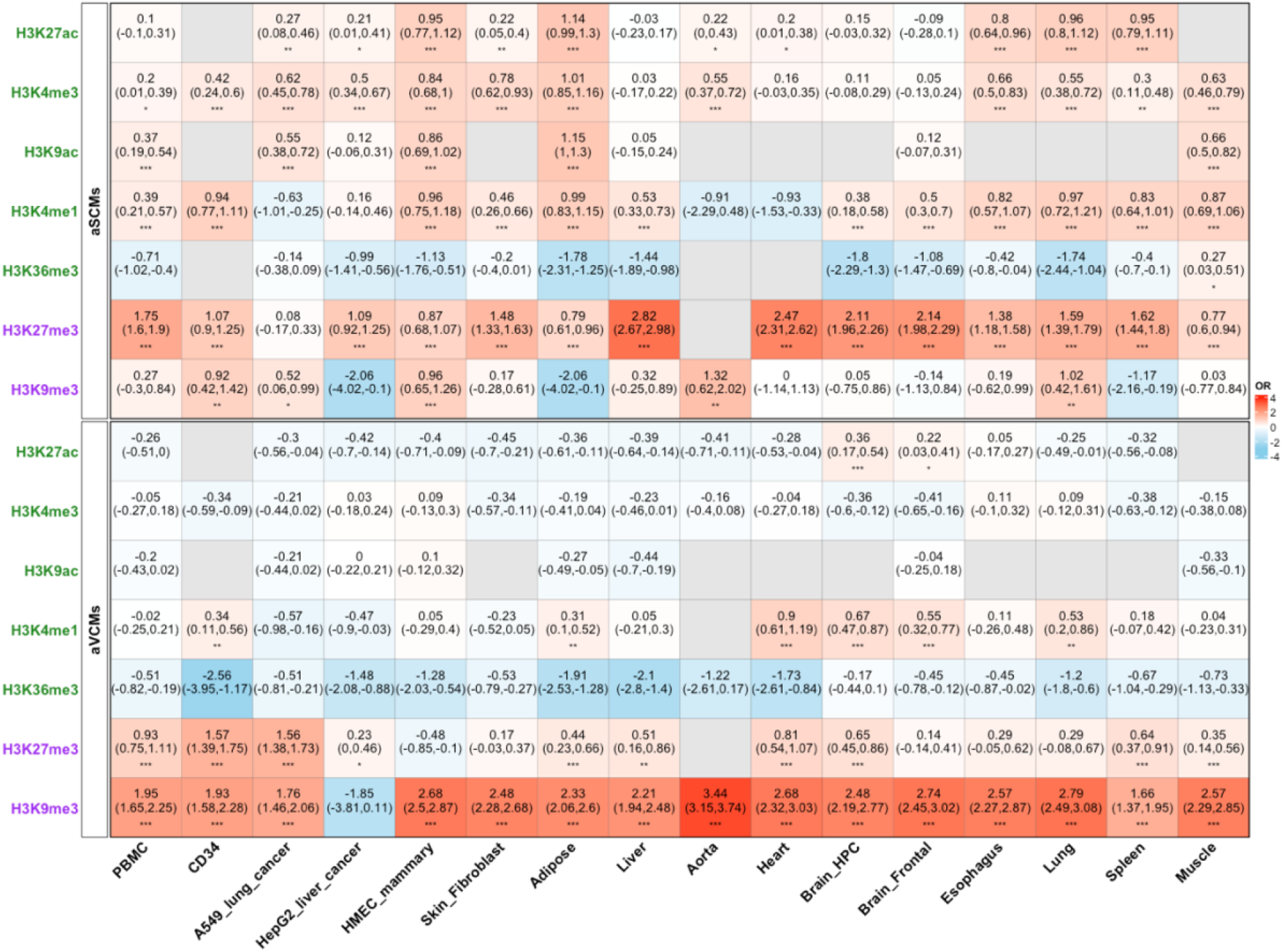
Enrichment of aSCMs (top) and aVCMs (bottom) for histone modifications across different cell lines/tissues. CpG modules identified via WGCNA^22^ were assessed for enrichment of histone marks across ENCODE cell lines. Overlap enrichment was assessed using one-sided hypergeometric test. The positive and negative log(OR) values represent over- and under-representation, respectively. The grey cells indicate absence of overlap. The values in brackets represent the confidence intervals for the OR (in log scale) and the asterisks are the significance strength (* = p < 0.05; ** = p < 0.01; *** = p < 0.001). The histone marks are grouped as active (green) and repressive (purple) marks. Abbreviations: age-stable co-methylation modules (aSCMs), age-variable co-methylation modules (aVCMs).

### Functional and phenotypic relevance of co-methylation modules

We used the region-based GREAT tool ^29^ to characterise the functional profiles of age-related co-methylation CpG sets. Both aSCMs and aVCMs were significantly enriched for developmental pathways (Fig. 6; Table S15a), although they mapped to distinct developmental processes. aSCMs were enriched for genes involved in developmental patterning and cellular identity, whereas aVCMs were associated with growth factor signalling and tissue morphogenesis (Fig. 6a,b), consistent with observations from aVCPs features (Fig. S2d).

**Figure 1.**
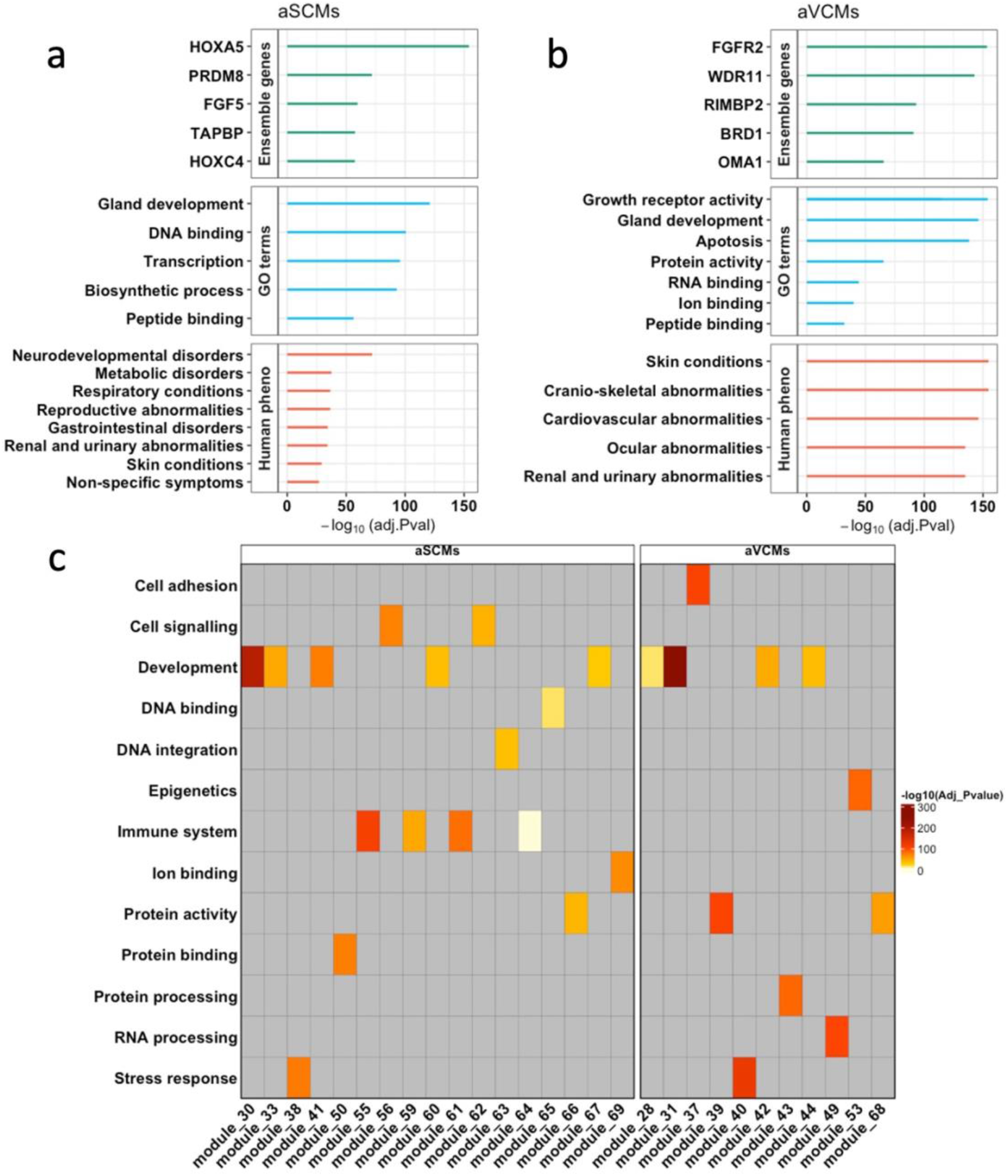
Biological themes associated with age-stable and age-variable co-methylation modules. **(a, b)** Enriched biological themes among combined aSCMs (**a**) and combined aVCMs (**b**). **(c)** Module-level distribution of enriched biological themes. Full enrichment results are provided in Supplementary Table 15a,b. Biological themes were manually assigned based on the top-ranked significantly enriched Gene Ontology (GO) terms for each module. Complete functional enrichment results for pooled and individual module analyses are provided in Supplementary Table S15a. Grey cells indicate that no significant enrichment was identified for the corresponding biological theme. Abbreviations: age-stable co-methylation modules (aSCMs), age-variable co-methylation modules (aVCMs).

Functional enrichment was also performed separately for each module to assess biological heterogeneity across modules. Distinct biological themes emerged across modules (Fig. 6c; Table S15a). Developmental pathways were enriched in several modules within both aSCMs and aVCMs, whereas immune-related enrichment was restricted to four aSCMs (modules 55, 59, 61 and 64). Stress-response pathways were shared across both module types. The findings also showed that aSCMs are enriched for neuro-psychiatric phenotypes, including mutism, psychosis and paranoia (Table S15a).

We next evaluated whether genes linked to aSCMs and aVCMs co-methylation features exhibit differential enrichment across age-related phenotypes and complex traits. Both co-methylation architectures showed nominal (p < 0.05) enrichment across various GWAS and MWAS gene hits; however, aSCMs were associated with cardiovascular and longevity-related traits, whereas aVCMs features showed stronger enrichment for hormonally regulated phenotypes (e.g., age at menarche, testosterone levels) (Fig. 7; Tables S16–S17; Supplementary Results). We observed limited overlap with curated transcriptomic aging gene sets or genes associated with longevity interventions (Fig. 7b and Fig. S17; Table S18; Supplementary Results). However, we found enrichment for second-generation epigenetic clocks (DunedinPACE and Pan-mammalian Clock 3; p < 0.01) but not first-generation epigenetic clocks. Nonetheless, these overlaps should be interpreted cautiously given the predictor-selection procedures used to construct most epigenetic clocks.

**Figure 1.**
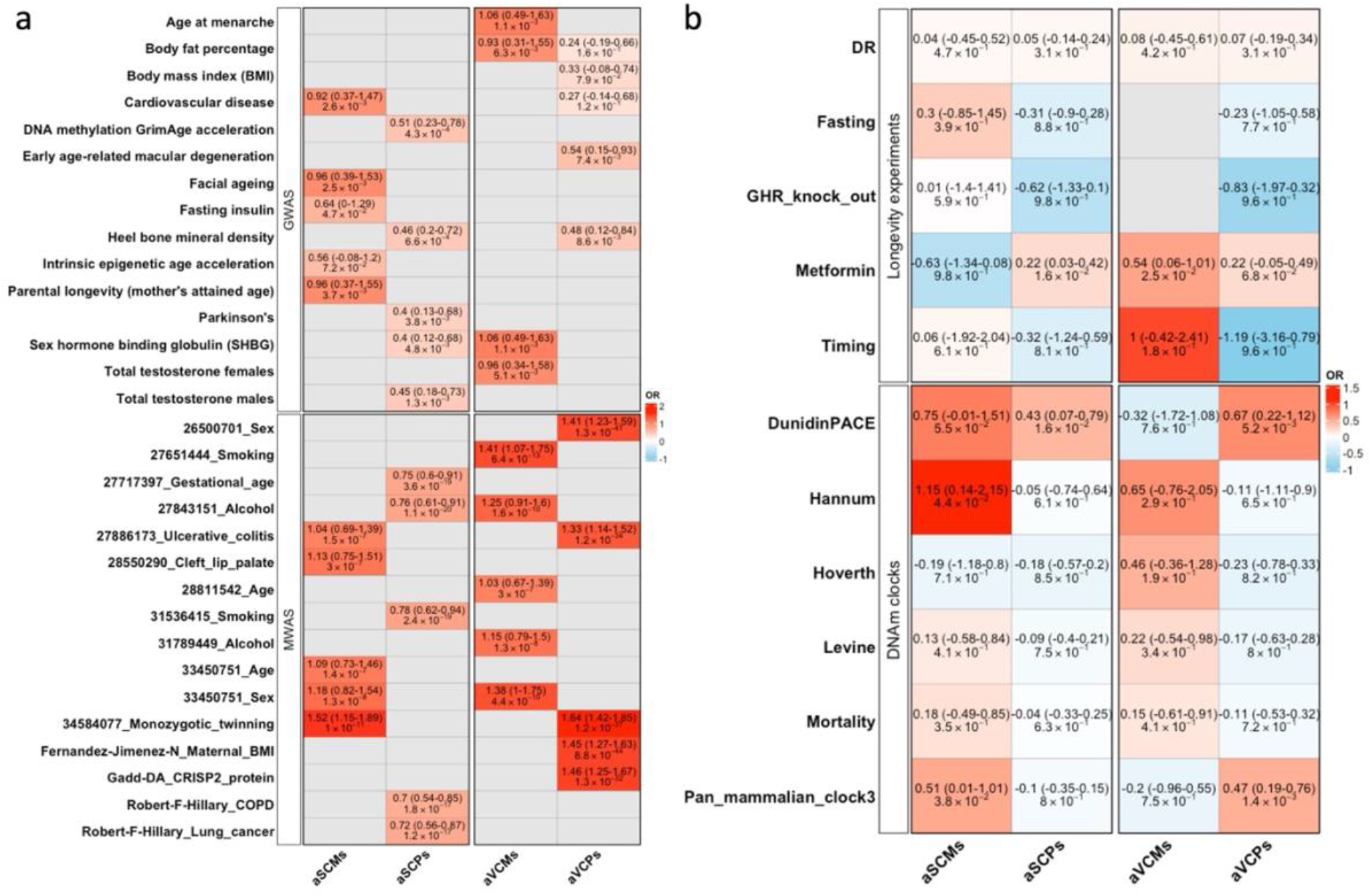
Enrichment of age-stable and age-variable co-methylation features among aging-related traits, longevity interventions, and epigenetic clocks.**(a)** Overlap between genes linked to age-stable and age-variable co-methylation features and genes identified from published GWAS and MWAS of complex traits and aging-related phenotypes. Numeric prefixes preceding trait names denote unique EWAS study identifiers for independent studies of the same phenotype. Grey cells indicate absence of overlap. **(b)** Overlap between age-related co-methylation features and genes associated with longevity interventions, as well as genes linked to CpGs incorporated in epigenetic clocks. Heat maps show odds ratios (Ors, in log scale) from hypergeometric enrichment tests. Values in parentheses indicate 95% confidence intervals for the ORs. Complete enrichment statistics are provided in Supplementary Table 16, 17 and 18. Grey cells indicate absence of overlap. Abbreviations: age-stable co-methylation modules (aSCMs), age-variable co-methylation modules (aVCMs), age-stable co-methylated probe pairs (aSCPs), age-variable co-methylated probe pairs (aVCPs), dietary restriction (DR), and growth hormone receptor (GHR), genome-wide association studies (GWAS), methylome-wide association studies (MWAS).

### Sex-stratified co-methylation analysis

Sex differences in age-associated DNA methylation dynamics are well documented in humans and other species ^30,31^. In addition, males and females exhibit differential rates of biological aging as captured by DNA methylation–based epigenetic clocks ^32^. We examined whether age-related co-methylation architecture differs between sexes by comparing module preservation changes between extreme age groups in males and females.

Preservation change scores were moderately correlated between sexes (r = 0.62), suggesting both shared and sex-specific patterns of network remodelling (Fig. S21). Using female- and male-specific reference modules derived from youngest age-group (Age1), we identified a subset of modules consistently showing sex-specific age-related disintegration. This includes correlated CpGs within protocadherin-β (*PCDHB*) gene cluster, which showed relatively stronger age-related remodelling in females than males (Fig. S21).

Notably, divergence in preservation statistics was most pronounced around midlife (50–55 years; Fig. S18), coinciding with the typical age of the menopausal transition in women. Functional enrichment analyses indicated that sex-biased modules are enriched for pathways related to immune regulation, metabolic processes, developmental signalling, and imprinting loci (see Supplementary Results for details; Fig. S21-22).

## Discussion

Through the analysis of genome-wide DNA methylation patterns across a large human cohort, we identified a spectrum of age-related patterns, ranging from modules that remain highly preserved (age-stable) across the lifespan to those that undergo progressive remodelling (age-variable) with age. Age-stable modules are associated with several features of genomic regulation, including trans-meQTLs, chromatin loops, TAD-associated organisation, and Polycomb-related regulatory elements. Genes linked to age-stable modules are enriched for immune pathways, whereas those associated with age-variable modules map to developmental pathways and hormonally regulated phenotypes. These findings extend our understanding of age-related DNA methylation change beyond shifts in mean levels and variance at individual CpG sites ^7,12,30,33-35^, towards a framework in which coordinated multi-site regulatory modules show distinct trajectories of stability and remodelling during aging.

Previous studies have shown that CpG sites exhibiting age-associated variability are enriched in repressive chromatin domains and depleted from active regulatory elements ^7,34,36^. In agreement, age-variable co-methylation modules from our work showed weaker enrichment for active regulatory regions, whereas age-stable modules were preferentially enriched within promoters, CpG islands and transcription start sites, suggesting that age-stable modules are linked to regions under stronger regulatory constraint. In support of these findings, we also observed that age-stable co-methylation modules were strongly enriched for trans-meQTLs, whereas age-variable modules showed a more balanced contribution of cis- and trans-genetic influences, indicating contrasting patterns of long-range genetic regulation. In addition, CpGs linked to age-stable modules are located within chromatin loops and topologically associating domain (TAD) boundaries)—genomic structures that facilitate distal regulatory interactions ^37^. Similarly, at the level of chromatin state, age-stable and age-variable modules also showed a clear divergence in regulatory context. Age-stable modules localise to regions marked by active regulatory features (H3K4me1 and H3K4me3) and Polycomb-associated repression (H3K27me3), whereas age-variable modules are enriched in H3K9me3-marked constitutive heterochromatin, consistent with previous reports e.g., ^7,34^. Notably, Polycomb-regulated chromatin maintains developmental genes in a constrained but reversible ‘poised’ state, whereas H3K9me3 marks compact, transcriptionally inert heterochromatin ^38^. The enrichment of age-variable modules within constitutive heterochromatin is consistent with evidence that age-related erosion of H3K9me3-marked heterochromatin contributes to disruption of chromatin organisation during aging ^39,40^.

Notably, our pairwise correlation results (Fig. 2) and previous longitudinal studies e.g., ^35^ have demonstrated that DNA methylation undergoes widespread and structured changes during human aging, affecting a substantial proportion of genome-wide CpG sites. However, our network-level results suggest that such extensive CpG-mean-level changes do not necessarily translate into age-related perturbation of higher-order CpG organisation.

Recent work ^41^ has proposed a systems-level view of aging as a progressive loss of epigenetic fidelity driven by interconnected failures in nuclear architecture, Polycomb-mediated memory, nucleosome composition and transcriptional regulation. Whereas our findings support this model, they further indicate that loss of epigenetic fidelity is heterogeneous across the methylome, with most co-methylation modules retaining stable organisation and only a subset exhibiting progressive age-related disruption.

Our study has several limitations. First, the study cohort consists predominantly of individuals of European ancestry, which may limit the generalisability of age-related co-methylation patterns across genetically diverse populations. Second, our analyses do not address co-methylation module dynamics during childhood or early adolescence because the study cohort comprised individuals aged 17–99 years. Third, the DNA methylation data were generated from bulk whole blood. Future studies incorporating additional tissues and single-cell methylation profiling will help determine the tissue specificity of age-related co-methylation features. Fourth, our results were derived from a cross-sectional cohort, which limits our ability to infer within-individual temporal dynamics of co-methylation. Although an independent external cohort was not available for replication, the consistent association of age-stable and age-variable modules across multiple independent genomic and regulatory annotations supports the robustness of our findings. Future work incorporating longitudinal and disease-specific cohorts will be important to determine how disruption of co-methylation modules relates to phenotypic heterogeneity and disease progression. Finally, experimental and integrative multi-omic approaches will be required to dissect the causal role of long-range genetic regulation and three-dimensional chromatin organisation in maintaining co-methylation module stability during aging and disease.

## Methods

### Generation Scotland cohort

Generation Scotland (GS), is a large family-structured cohort study that consists of more than 24,000 individuals from across Scotland, as described previously ^20,21^. Participants were identified via Community Health Index numbers, with the support of Scottish Practices and Professionals Involved in Research. The initial phase of recruitment (2006 to 2011) focussed on the Glasgow and Tayside regions of Scotland and was later extended to Ayrshire, Arran, and the Northeast of Scotland. Detailed health and lifestyle information were collected via questionnaires at the study baseline alongside venepuncture to obtain whole blood samples from which DNAm was assayed.

### Data filtering, pre-correction and age group definition

Participants in the GS cohort were assayed for over 850,000 CpG sites. In this study, we filtered out probes that targeted polymorphic CpG sites (i.e., CpG sites where a SNP occurs at the cytosine or guanine position, resulting in the loss or gain of a CpG site) or those shown to cross-hybridize with multiple regions of the genome. After removing related individuals (genetic relationship matrix [GRM] < 0.05), the final DNA methylation dataset consisted of 7,532 individuals and 752,722 probes. This dataset was pre-corrected for estimated cell proportions (Bcell, CD4T, CD8T, Eos, Mono, Neu and NK) and other covariates, including age, sex, batch effects, and smoking status, as follows:

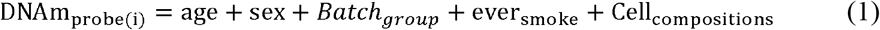

Age-associated effects on mean DNAm levels were removed prior to analysis to ensure that downstream network analyses captured age-related changes in correlation structure rather than age-related shifts in average methylation levels. The pre-corrected DNA methylation dataset was divided into eight similarly sized age groups (952–986 individuals per group), ensuring comparable statistical precision for co-methylation analyses across age strata (Fig. 1 and Fig. S1). The distribution of individuals across the age groups is shown in Figure S1.

### Statistical analysis

#### Pairwise correlation (co-methylation) analysis

Co-methylated CpG pairs were identified through pairwise correlations of DNA methylation probes within each chromosome across age groups (Age groups 1–8). Strongly co-methylated probe pairs were defined as those with absolute correlation coefficients exceeding 0.8 (|r| > 0.8) in at least age group. These probe pairs were then tracked across all age groups to assess changes in co-methylation (correlation) strength.

To characterise local co-methylation structure, we examined the relationship between correlation strength and genomic distance (≤10 kb), separately for positively and negatively correlated probe pairs.

Probe pairs exhibiting age-dependent changes in co-methylation variability were identified using Breusch–Pagan test of heteroskedasticity. Probe pairs showing significant increases in variance across age groups were classified as age-variable co-methylated probe pairs (aVCPs), whereas those with minimal change in variance were defined as age-stable co-methylated probe pairs (aSCPs). Detailed statistical analyses and model specifications are provided in the Supplementary Methods.

#### Network co-methylation analysis and preservation

Age-related co-methylation features at the network level were identified using Weighted Correlation Network Analysis (WGCNA) to construct modules of co-methylated CpG sites and evaluate their preservation across age groups ^22,23^. Signed modules were constructed using the *blockwiseModules* function with biweight midcorrelation (bicor) and a soft-thresholding power of 6, selected to approximate scale-free topology. Modules were detected using a minimum module size of 20, with *deepSplit* = 3 and *mergeCutHeight* = 0.10. The youngest age group (<36 years) was used as the reference dataset, and modules identified in this group were subsequently assessed for preservation across the remaining age groups (Age groups 2–8), each treated as a test dataset. Preservation was evaluated using both density-based metrics (proportion of variance explained) and connectivity-based statistics (cor.cor, cor.kIM, cor.kME).

Age-related trends in co-methylation modules were characterised by modelling preservation statistics as a function of age using Generalized Additive Models, allowing for non-linear trajectories across age groups. Modules showing significant age-associated trends (p < 0.05) were classified as candidate age-variable modules, whereas non-significant modules were classified as candidate age-stable modules. This analysis was performed separately for each WGCNA preservation metric.

Next, we quantified the percentage change in preservation statistics between the youngest and oldest age groups (Age group 1 vs. Age group 8) as follows:

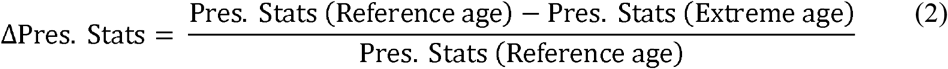

Candidate age-variable modules were retained only if they exceeded a minimum change threshold, defined as the greater of the 50th percentile of the preservation-change distribution and a metric-specific fixed threshold: >5% for cor.kME and >10% for proportion of variance explained. Visual inspection of preservation trajectories indicated that these thresholds provided good separation between modules showing stable and changing preservation patterns across age groups. Modules with unstable or highly variable trajectories across age groups (standard deviation above the 40th percentile of the observed distribution), or failing to meet the above criteria, were not assigned to either category. In addition, modules with low connectivity in the youngest test age group (preservation < 0.40 in Age group 2) were also not assigned to either category. The remaining modules were classified as age-variable (aVCMs) and age-stable (aSCMs) co-methylation modules for downstream analyses. The workflow of module selection criteria is illustrated in Supplementary Fig. S9.

#### Genetic and environmental (smoke) related influences

To investigate association of the selected co-methylation CpG sets with genetic effects we performed one-sided hypergeometric overlap test with methylation quantitative trait loci ^9^. For each cis-and trans meQTLs, we selected top 10,000 hits corresponding to a p-value of 8.10 × 10 ^−131^ and 9.11 × 10 ^−11^, respectively. The background set for the analysis consisted of 752,722 CpG sites—the total number of CpGs retained after quality control.

To evaluate smoke-related environmental contributions, we used omics-based restricted maximum likelihood (OREML) implemented in OSCA ^42^ to estimate the proportion of variance in smoking-related phenotypes (ever-smoked status and pack-years) explained by each co-methylation CpG set. Models included age, sex, batch effects, and cell-type composition as covariates. Full model specifications and matrix formulations are provided in the Supplementary Methods.

#### Histone marks

We obtained histone modification data from the Roadmap Epigenomics Project ^28^. Seven core histone marks (H3K4me1, H3K4me3, H3K9ac, H3K9me3, H3K27ac, H3K27me3, and H3K36me3) were selected for overlap analysis across 16 tissues. The top 10,000 ranked histone mark peaks for each tissue, following Lu, et al. ^34^, were selected and their genomic coordinates intersected with the study CpG sets. Enrichment of the age-related CpG sets (aSCPs/aVCPs, aSCMs/aVCMs) within histone marks was evaluated using a one-sided hypergeometric test for each tissue, separately. The background set included 752,722 CpG sites.

#### Three-dimensional chromatin organization

We obtained Hi-C chromatin loop and topologically associating domain (TAD) coordinates from the 3D Genome Browser (https://3dgenome.fsm.northwestern.edu; accessed February 2025). Chromatin interaction data were derived from the ENCODE Phase III project and included 11 human cell lines. To define TAD boundary regions, we extended 100 kb upstream and downstream of each TAD start and end coordinate, respectively. We then overlapped our age-related CpG sets with chromatin loop anchors and TAD boundary regions and assessed enrichment using a one-sided hypergeometric test, with all quality-controlled CpGs (N = 752,722) serving as the background set.

#### Functional enrichment analysis

The probes within age-related modules (aSCMs/aVCMs) were used as the foreground input set for hypergeometric analysis using GREAT software ^43^. We applied the ‘basal plus extension’ rule, where genomic regions (probe locations) were associated with nearby genes. Each gene was assigned a basal regulatory domain with a minimum distance of 5 kb upstream and 1 kb downstream of the transcription start site (TSS), regardless of other nearby genes. This basal domain was extended up to 1,000 kb in both directions, but no further than the maximum extension toward the nearest gene’s basal domain. The entire probe set (N = 752,722) from our study was used as the background set in the GREAT hypergeometric test.

The age-related co-methylated probe pairs identified from the heteroskedasticity analysis (aSCPs and aVCPs) were similarly annotated using GREAT.

#### EWAS overlap analysis

We assessed whether genes linked to age-related co-methylation signatures were enriched among genes identified in published epigenome-wide association studies (EWAS). EWAS summary statistics were obtained from the EWAS Catalog (http://www.ewascatalog.org). For each study or model, analyses were restricted to autosomal CpGs, which were ranked by statistical significance and the top 1,000 CpGs were selected to yield a consistent number of high-confidence associations across studies with differing sample sizes and statistical power.

Selected CpGs were mapped to putative target genes using the basal plus extension association rule implemented in GREAT ^43^. EWAS studies mapping to fewer than 100 genes were excluded to avoid overlap analyses based on sparse gene sets, leaving 1,990 studies.

For each co-methylation set (aSCPs, aVCPs, aSCMs, aVCMs), enrichment was assessed using one-sided hypergeometric tests. The background consisted of 17,786 unique genes associated with all autosomal EWAS CpGs passing selection criteria.

#### Overlap with GWAS traits

We assessed the enrichment of genes linked to age-related CpG sets in genome-wide association study (GWAS) traits, following the approach of Lu, et al. ^34^. GWAS summary statistics from 89 studies covering age-related, anthropometric, hormonal, longevity, and disease traits were curated (Table S23).

CpGs were mapped to their nearest genes (within 5 kb upstream and 1 kb downstream; Build 37), and SNPs from each GWAS dataset were mapped to genes within a ±50 kb window. Gene-level association statistics were computed using MAGMA v1.10 ^44^ with a European reference panel from the 1000 Genomes Project ^45^.

For each GWAS, the top 2.5% and 5% of genes based on gene-level p-values were selected and tested for overlap with genes linked to aSCPs/aVCPs and aSCMs/aVCMs using one-sided hypergeometric tests. The background consisted of 17,593 genes associated with all CpGs retained after quality control (N = 752,722). To avoid bias from the major histocompatibility complex (MHC) region, genes located within the MHC region (chromosome 6; 25–34 Mb) were excluded before analysis.

## Supporting information

Supplementary Results

## Acknowledgements

Generation Scotland received core support from the Chief Scientist Office of the Scottish Government Health Directorates [CZD/16/6] and the Scottish Funding Council [HR03006] and is currently supported by the Wellcome Trust [216767/Z/19/Z]. Genotyping of the GS:SFHS samples were carried out by the Genetics Core Laboratory at the Edinburgh Clinical Research Facility, University of Edinburgh, Scotland and was funded by the Medical Research Council UK and the Wellcome Trust (Wellcome Trust Strategic Award “STratifying Resilience and Depression Longitudinally” (STRADL) Reference 104036/Z/14/Z). R.E.M and A.F.M are supported by the Biotechnology and Biological Sciences Research Council grant UKRI/BB/C001941/1.

## Data and code availability

The Generation Scotland data underlying this study are available through the Generation Scotland Access Committee upon reasonable request and subject to the relevant approvals. Information on the data access process is available at https://www.ed.ac.uk/generation-scotland/for-researchers. Source data supporting the findings of this study are provided within the article and its Supplementary Information where appropriate. Analysis scripts and code used to generate the results and figures will be deposited in a public repository before publication.

## Ethics approval

This study involves human participants. All components of Generation Scotland received ethical approval from the NHS Tayside Committee on Medical Research Ethics (REC reference: 05/S1401/89). Generation Scotland has also been granted Research Tissue Bank status by the East of Scotland Research Ethics Service (REC reference: 20/ES/0021), providing ethical approval for a broad range of medical research. Written informed consent was obtained from all GS:SFHS participants, and Next Generation Scotland (NGS) participants provided informed consent electronically prior to participation.

## Author contributions

A.F.M. conceived and designed the study. E.K.C, A.A.H and A.F.M. contributed to the formal data analysis. E.K.C. wrote the first draft. D.L.M. contributed to the acquisition and curation of the Generation Scotland data resource. R.E.M. contributed to the design and oversight of the Generation Scotland resource and revised the manuscript. All authors reviewed and approved the final manuscript for publication.

## References

1 Jones, M. J., Goodman, S. J. & Kobor, M. S. DNA methylation and healthy human aging. Aging Cell 14, 924–932 (2015). 10.1111/acel.12349

2 López-Otín, C., Blasco, M. A., Partridge, L., Serrano, M. & Kroemer, G. Hallmarks of aging: An expanding universe. Cell 186, 243–278 (2023). 10.1016/j.cell.2022.11.001

3 Levine, M. E. et al. Menopause accelerates biological aging. Proc Natl Acad Sci U S A 113, 9327–9332 (2016). 10.1073/pnas.1604558113

4 Bernabeu, E. et al. Refining epigenetic prediction of chronological and biological age. Genome Med 15, 12 (2023). 10.1186/s13073-023-01161-y

5 Horvath, S. DNA methylation age of human tissues and cell types. Genome Biol 14, R115 (2013). 10.1186/gb-2013-14-10-r115

6 Levine, M. E. et al. An epigenetic biomarker of aging for lifespan and healthspan. Aging (albany NY) 10, 573 (2018).

7 Slieker, R. C. et al. Age-related accrual of methylomic variability is linked to fundamental ageing mechanisms. Genome Biol 17, 191 (2016). 10.1186/s13059-016-1053-6

8 Banovich, N. E. et al. Methylation QTLs are associated with coordinated changes in transcription factor binding, histone modifications, and gene expression levels. PLoS Genet 10, e1004663 (2014). 10.1371/journal.pgen.1004663

9 Min, J. L. et al. Genomic and phenotypic insights from an atlas of genetic effects on DNA methylation. Nat Genet 53, 1311–1321 (2021). 10.1038/s41588-021-00923-x

10 Guo, S. et al. Identification of methylation haplotype blocks aids in deconvolution of heterogeneous tissue samples and tumor tissue-of-origin mapping from plasma DNA. Nat Genet 49, 635–642 (2017). 10.1038/ng.3805

11 Ochana, B.-L. et al. Time is encoded by methylation changes at clustered CpG sites. Cell Reports 44, 115958 (2025). 10.1016/j.celrep.2025.115958

12 Haghani, A. et al. DNA methylation networks underlying mammalian traits. Science 381, eabq5693 (2023). 10.1126/science.abq5693

13 Harvey, J. et al. Epigenetic insights into neuropsychiatric and cognitive symptoms in Parkinson’s disease: A DNA co-methylation network analysis. NPJ Parkinsons Dis 11, 39 (2025). 10.1038/s41531-025-00877-5

14 Kim, J. P. et al. Integrative Co-methylation Network Analysis Identifies Novel DNA Methylation Signatures and Their Target Genes in Alzheimer’s Disease. Biol Psychiatry 93, 842–851 (2023). 10.1016/j.biopsych.2022.06.020

15 Kouhsar, M. et al. A brain DNA co-methylation network analysis of psychosis in Alzheimer’s disease. Alzheimers Dement 21, e14501 (2025). 10.1002/alz.14501

16 Horvath, S. et al. Aging effects on DNA methylation modules in human brain and blood tissue. Genome Biol 13, R97 (2012). 10.1186/gb-2012-13-10-r97

17 Jacques, M. et al. DNA Methylation Ageing Atlas Across 17 Human Tissues. bioRxiv, 2025.2007. 2021.665830 (2025).

18 Watkins, S. H. et al. DNA co-methylation has a stable structure and is related to specific aspects of genome regulation. bioRxiv, 2022.2003. 2016.484648 (2022).

19 Jensen, D. et al. Co-methylation networks associated with cognition and structural brain development during adolescence. Front Genet 15, 1451150 (2024). 10.3389/fgene.2024.1451150

20 Walker, R. M. et al. Data Resource Profile: Whole-Blood DNA Methylation Resource in Generation Scotland (MeGS). Int J Epidemiol 54 (2025). 10.1093/ije/dyaf091

21 Milbourn, H. et al. Generation Scotland: an update on Scotland’s longitudinal family health study. BMJ Open 14, e084719 (2024). 10.1136/bmjopen-2024-084719

22 Langfelder, P. & Horvath, S. WGCNA: an R package for weighted correlation network analysis. BMC Bioinformatics 9, 559 (2008). 10.1186/1471-2105-9-559

23 Langfelder, P., Luo, R., Oldham, M. C. & Horvath, S. Is my network module preserved and reproducible? PLoS Comput Biol 7, e1001057 (2011). 10.1371/journal.pcbi.1001057

24 Szklarczyk, D. et al. The STRING database in 2021: customizable protein-protein networks, and functional characterization of user-uploaded gene/measurement sets. Nucleic Acids Res 49, D605–d612 (2021). 10.1093/nar/gkaa1074

25 Vu, H. & Ernst, J. Universal annotation of the human genome through integration of over a thousand epigenomic datasets. Genome Biol 23, 9 (2022). 10.1186/s13059-021-02572-z

26 Wang, Y. et al. The 3D Genome Browser: a web-based browser for visualizing 3D genome organization and long-range chromatin interactions. Genome biology 19, 1–12 (2018).

27 Davis, C. A. et al. The Encyclopedia of DNA elements (ENCODE): data portal update. Nucleic acids research 46, D794–D801 (2018).

28 Roadmap Epigenomics Consortium. Integrative analysis of 111 reference human epigenomes. Nature 518, 317–330 (2015).

29 McLean, C. Y. et al. GREAT improves functional interpretation of cis-regulatory regions. Nature biotechnology 28, 495–501 (2010).

30 Yusipov, I. et al. Age-related DNA methylation changes are sex-specific: a comprehensive assessment. Aging (Albany NY) 12, 24057–24080 (2020). 10.18632/aging.202251

31 Shealy, E. P., Schwartz, T. S., Cox, R. M., Reedy, A. M. & Parrott, B. B. DNA methylation-based age prediction and sex-specific epigenetic aging in a lizard with female-biased longevity. Sci Adv 11, eadq3589 (2025). 10.1126/sciadv.adq3589

32 Crimmins, E. M., Thyagarajan, B., Levine, M. E., Weir, D. R. & Faul, J. Associations of Age, Sex, Race/Ethnicity, and Education With 13 Epigenetic Clocks in a Nationally Representative U.S. Sample: The Health and Retirement Study. J Gerontol A Biol Sci Med Sci 76, 1117–1123 (2021). 10.1093/gerona/glab016

33 Seale, K., Teschendorff, A., Reiner, A. P., Voisin, S. & Eynon, N. A comprehensive map of the aging blood methylome in humans. Genome Biol 25, 240 (2024). 10.1186/s13059-024-03381-w

34 Lu, A. T. et al. Universal DNA methylation age across mammalian tissues. Nat Aging 3, 1144–1166 (2023). 10.1038/s43587-023-00462-6

35 Mulder, R. H. et al. Epigenome-wide change and variation in DNA methylation in childhood: trajectories from birth to late adolescence. Hum Mol Genet 30, 119–134 (2021). 10.1093/hmg/ddaa280

36 Wang, Y. et al. Epigenetic influences on aging: a longitudinal genome-wide methylation study in old Swedish twins. Epigenetics 13, 975–987 (2018). 10.1080/15592294.2018.1526028

37 Batut, P. J. et al. Genome organization controls transcriptional dynamics during development. Science 375, 566–570 (2022). 10.1126/science.abi7178

38 Schuettengruber, B., Bourbon, H. M., Di Croce, L. & Cavalli, G. Genome Regulation by Polycomb and Trithorax: 70 Years and Counting. Cell 171, 34–57 (2017). 10.1016/j.cell.2017.08.002

39 Mrabti, C. et al. Loss of H3K9 trimethylation leads to premature aging. bioRxiv (2024). 10.1101/2024.07.24.604929

40 Tsurumi, A. & Li, W. X. Global heterochromatin loss: a unifying theory of aging? Epigenetics 7, 680–688 (2012). 10.4161/epi.20540

41 Yücel, A. D. & Gladyshev, V. N. Systemic epigenetic dysregulation as a driver of ageing and a therapeutic target. Nat Rev Mol Cell Biol (2026). 10.1038/s41580-026-00958-0

42 Zhang, F. et al. OSCA: a tool for omic-data-based complex trait analysis. Genome biology 20, 1–13 (2019).

43 Gu, Z. & Hübschmann, D. rGREAT: an R/bioconductor package for functional enrichment on genomic regions. Bioinformatics 39 (2023). 10.1093/bioinformatics/btac745

44 de Leeuw, C. A., Mooij, J. M., Heskes, T. & Posthuma, D. MAGMA: generalized geneset analysis of GWAS data. PLoS computational biology 11, e1004219 (2015).

45 1000 Genomes Project Consortium. A global reference for human genetic variation. Nature 526, 68 (2015).

46 Yang, J., Lee, S. H., Goddard, M. E. & Visscher, P. M. GCTA: a tool for genome-wide complex trait analysis. The American Journal of Human Genetics 88, 76–82 (2011).

47 Zhou, W. et al. DNA methylation loss in late-replicating domains is linked to mitotic cell division. Nature genetics 50, 591–602 (2018).

