## Supplementary Methods for "Age-related dynamics of DNA co-methylation modules in humans"

### Pairwise correlation analysis across age groups

To identify co-methylated CpG probe pairs, we calculated pairwise Pearson correlations of DNA methylation levels for all probes within each chromosome, separately for each age group. From these results, we selected strongly co-methylated probe pairs in each reference age group (Age groups 1–8), defined as those with correlation coefficients exceeding |r| > 0.8 (i.e., r > 0.8 or r < –0.8).

The selected co-methylated probe pairs were then carried forward and their pairwise correlations recalculated across all remaining age groups to evaluate changes in co-methylation strength with age. To minimise bias associated with reference group selection, this procedure was repeated iteratively, using each age group (Age groups 1–8) as the reference for defining strongly co-methylated probe pairs.

### Distance-dependent co-methylation patterns

To examine how co-methylation varies with genomic distance, we restricted the analysis to probe pairs located within 10 kb of each other. Probe pairs were separated into positively (r > 0.8) and negatively (r < –0.8) correlated sets.

For each subset, probe pairs were binned into 200 groups based on genomic distance. Within each bin, we calculated the mean pairwise distance and the corresponding mean correlation coefficient, along with standard deviations. These summaries were used to characterise the relationship between genomic proximity and co-methylation strength.

### Heteroskedasticity analysis of pairwise co-methylation dynamics

To identify probe pairs showing age-dependent changes in co-methylation variability, we applied a Breusch–Pagan test of heteroskedasticity ^1^, using a two-stage modelling framework.

First, we modelled the relationship between paired co-methylated probes while adjusting for age group:

$${probe}_{1}={probe}_{2}+Age_{group}+error$$

(1)

Residuals from this model were then used to assess variance changes across age groups. Specifically, squared residuals were normalised by their mean and regressed against age group:

$${normalized}_{{residuals}^{2}}=Age_{group}+error$$

(2)

Under the null hypothesis of homoscedasticity (equal variance across age groups), test statistics follow a chi-squared distribution.

Probe pairs were ranked based on the significance of the heteroskedasticity test. The top 1,000 probe pairs with the smallest p-values were classified as age-variable co-methylation probe pairs (aVCPs), representing those with increasing variability across age. Conversely, the bottom 1,000 probe pairs with p-values ~1.0 were defined as age-stable co-methylation probe pairs (aSCPs).

### Cross-study overlap analysis

We assessed the overlap between genes linked to age-related co-methylation features and age-variable methylation positions reported by Slieker, et al. ^2^. Overlap significance was evaluated using a one-sided hypergeometric test, with either all CpGs retained after quality control or genes mapped to these CpGs used as the background, as appropriate.

### Genetic and environmental contributions

To assess environmental contributions, we used restricted maximum likelihood (OREML) based on OSCA (OmicS-data-based Complex trait Analysis) software ^3^ to calculate the proportion of variance in smoking phenotypes (ever-smoked status and pack-years) explained by the selected co-methylation CpG sets. The OREML model as follows:

$$y=\mathrm{Cb}+\mathrm{Wu}+e$$

(3)

where $\mathbf{y}$ is an $n\times1$ vector of smoke phenotypes; $n$ = number of samples; **C** = covariates (age, sex, batch effects, and cell compositions); **W** = $n\times m$matrix DNAm probes ($m$ = total number of DNAm for the selected CpG sets; Table S2 and S4); $\mathbf{b}$ is the vector of fixed effects; $\mathbf{u}$ is an $m\times1$vector of joint effects of all DNAm probes on smoking phenotypes [$\boldsymbol{u}$ ~$N(0, {\mathbf{I}\sigma}_{u}^{2})$]; $\boldsymbol{e}$ = vector of random residuals [$\boldsymbol{e}$ ~$N(0, {\mathbf{I}\sigma}_{e}^{2})$]. The variance (***y***) = **V** = $WW^{'}\sigma_{u}^{2}+ {\mathbf{I}\sigma}_{e}^{2}$ = $\mathbf{A}\sigma_{o}^{2}+ {\mathbf{I}\sigma}_{e}^{2}$ with **A** = $WW^{'}/m$ and $\sigma_{o}^{2}$ = ${m\sigma}_{u}^{2}$. **A** is the omics-data relationship matrix (ORM) computed as:

$$A_{jk}= \frac{1}{m}\sum_{i} \frac{(x_{ij}-u_{i})\left( x_{ik}-u_{i} \right)}{\sigma_{i}^{2}}$$

(4)

where $A_{jk}$is the relationship between individual *j* and *k*; $x_{ij}$is the unstandardized DNAm level of probe *i* in individual *j*; $u_{i}$ and $\sigma_{i}^{2}$ are the mean and variance of probe *i*, respectively*;* and $m$ is the number of DNAm probes for each selected CpG set. The proportion of variance ($R^{2}$) for ever-smoked status and pack-years captured by the selected CpG set was calculated as: $R^{2}= {\sigma_{o}^{2}}/{(\sigma_{o}^{2}+\sigma_{e}^{2}}$).

### Chromatin states

We conducted chromatin state enrichment analysis using the human chromatin annotation resource from Vu and Ernst ^4^. These authors profiled chromatin states for the human genome using over 1000 datasets from more that 100 cell types. We overlaid our age-related CpG sets within the chromatin state genomic bins and performed one-sided hypergeometric test to assess the degree of enrichment for each state. The background set for the analysis consisted of the total number of CpGs retained after quality control.

### Transcription factor binding sites

To determine whether the selected CpG sets were overrepresented within specific transcription factor binding sites (TFBS), we obtained TFBS data from the ENCODE database ^5^. TFBS were first overlapped with the background CpG set (N = 752,722), yielding 536,262 CpGs associated with 380 transcription factors (TFs). To reduce noise and focus on biologically relevant TFs, we retained only those linked to more than five but fewer than 2,000 genes. This filtering step resulted in 42 TFs corresponding to 48,880 CpGs, with the number of CpGs per TF ranging from 41 (RBM17) to 5,196 (CBX8).

### Chromatin accessibility

We downloaded age-related chromatin accessibility data from Patrick, et al. ^6^. These authors profiled differentially accessible chromatin regions (DARs) between young (2 months) and old (24 months) mice across 22 cell types. The DARs were classified into two groups: those that gain accessibility with age (opening DARs) and those that lose accessibility with age (closing DARs).

Before the overlap analysis, CpG coordinates (hg19) were converted to the mouse genome assembly (mm10) using the UCSC *liftOver* tool (<https://bioconductor.org/packages/liftOver>). For each tissue, ATAC-seq peaks were ranked by adjusted *p* values, and the top 10,000 peaks selected for analysis. We then tested whether our age-related CpG sets were over-represented in either opening or closing DARs within each tissue using a one-sided hypergeometric test. CpG sites retained after quality control (N = 752,722) served as the background set.

### Methylation domain architecture

We downloaded partially methylated domains (PMDs), highly methylated domains (HMDs), and solo-WCGW domain annotations from Zhou, et al. ^7^. Solo-WCGW sites are CpG dinucleotides flanked by ‘A:T’ bases on both sides or lacking neighbouring CpGs and are prone to hypomethylation during DNA replication. For each domain, we defined 100 kb windows upstream and downstream of the domain coordinates. Our age-related CpG sets were overlapped with these domains and enrichment was assessed using a one-sided hypergeometric test, with all CpGs that were retained after quality control used as the background set.

### Overlap with age-related phenotypes

We obtained human age-related gene data from several sources: the Digital Ageing Atlas ^8^, the Human and Musculus Aging Atlas ^9^, and Human Aging Genomic Resources ^10^. The genes from these resources were aggregated into a transcriptomic category. Additionally, we retrieved age-related variably methylated probes from Slieker, et al. ^2^ and genes associated with 2,000 age-related probes from the Mammalian Methylation Consortium ^11^, categorizing them into aVMPs and pan-mammalian categories, respectively. We also included genes associated with mortality risk from ^12^ and ^13^.

We performed a hypergeometric test, to assess the overlap between the selected genes located near or within age-related CpG sets. For sensitivity analysis, we generated a null distribution by randomly drawing 491 genes (matching the number of significant genes from the aVCPs-associated heteroskedasticity test) from the full gene set (N = 17,593) located near study probes. This random sampling was repeated 100 times to produce replicate sets and the overlap odds ratios.

### Sex-specific networks analysis

We performed sex-stratified network analysis to investigate the presence of sex-biased modules—that is, modules whose co-methylation structure is preserved with age in one sex but not in the other ^14,15^. To achieve this, we first pre-corrected the DNAm data as described in the main Methods, including all covariates in the model except for sex.

$$\mathrm{DNAm}_{probe(i)}=age+{Batch}_{group}+\mathrm{ever}_{\mathrm{smoke}}+\mathrm{Cell}_{\mathrm{compositions}}$$

(5)

The dataset was then divided by sex and further stratified into eight age groups within each sex, as described in Section 2.1. Co-methylation networks were constructed using the youngest age group from each sex (Age group 1: < 36 years), and their preservation statistics assessed across older age groups (Age groups 2–8) within the same sex as well as across the opposite sex. The number of samples in ‘Age group 1’ used for network construction in males and females was 433 and 543, respectively (Fig. S1). Network construction and module detection were performed using the same WGCNA framework and parameters described in the main Methods.

Sex-biased modules were defined as those showing a substantial decrease in network preservation estimates between the youngest and oldest age groups (i.e., Age groups 2 vs. 8) in one sex relative to the other. Finally, functional enrichment analyses were performed for the selected sex-biased modules.

### References

1	Breusch, T. S. & Pagan, A. R. A simple test for heteroscedasticity and random coefficient variation. *Econometrica: Journal of the econometric society*, 1287-1294 (1979).

2	Slieker, R. C. *et al.* Age-related accrual of methylomic variability is linked to fundamental ageing mechanisms. *Genome Biol* **17**, 191 (2016). <https://doi.org/10.1186/s13059-016-1053-6>

3	Zhang, F. *et al.* OSCA: a tool for omic-data-based complex trait analysis. *Genome biology* **20**, 1-13 (2019).

4	Vu, H. & Ernst, J. Universal annotation of the human genome through integration of over a thousand epigenomic datasets. *Genome Biol* **23**, 9 (2022). <https://doi.org/10.1186/s13059-021-02572-z>

5	Davis, C. A. *et al.* The Encyclopedia of DNA elements (ENCODE): data portal update. *Nucleic acids research* **46**, D794-D801 (2018).

6	Patrick, R. *et al.* The activity of early-life gene regulatory elements is hijacked in aging through pervasive AP-1-linked chromatin opening. *Cell Metab* **36**, 1858-1881.e1823 (2024). <https://doi.org/10.1016/j.cmet.2024.06.006>

7	Zhou, W. *et al.* DNA methylation loss in late-replicating domains is linked to mitotic cell division. *Nature genetics* **50**, 591-602 (2018).

8	Craig, T. *et al.* The Digital Ageing Atlas: integrating the diversity of age-related changes into a unified resource. *Nucleic acids research* **43**, D873-D878 (2015).

9	Aging Atlas Consortium. Aging Atlas: a multi-omics database for aging biology. *Nucleic acids research* **49**, D825-D830 (2021).

10	Tacutu, R. *et al.* Human ageing genomic resources: new and updated databases. *Nucleic acids research* **46**, D1083-D1090 (2018).

11	Lu, A. T. *et al.* Universal DNA methylation age across mammalian tissues. *Nat Aging* **3**, 1144-1166 (2023). <https://doi.org/10.1038/s43587-023-00462-6>

12	Levine, M. E. *et al.* An epigenetic biomarker of aging for lifespan and healthspan. *Aging (albany NY)* **10**, 573 (2018).

13	Zhang, Y. *et al.* DNA methylation signatures in peripheral blood strongly predict all-cause mortality. *Nature communications* **8**, 14617 (2017).

14	Langfelder, P. & Horvath, S. WGCNA: an R package for weighted correlation network analysis. *BMC Bioinformatics* **9**, 559 (2008). <https://doi.org/10.1186/1471-2105-9-559>

15	Langfelder, P., Luo, R., Oldham, M. C. & Horvath, S. Is my network module preserved and reproducible? *PLoS Comput Biol* **7**, e1001057 (2011). <https://doi.org/10.1371/journal.pcbi.1001057>
