## Supplementary Results for "Age-related dynamics of DNA co-methylation modules in humans"

### Characteristics of pairwise co-methylation relationships across age

Larger proportion of co-methylated probe pairs (selected across all age groups) were separated by large genomic distances (>10 Mb; 64.3%), whereas 20.5% occurred within 1 Mb and 15.2% were separated by 1–10 Mb (Fig. S5a). Co-methylation strength declined with increasing genomic distance and plateaued at approximately 1 kb (Fig. S5b). Among positively co-methylated probe pairs, correlation strength in the oldest age group (Age8) decreased modestly with increasing genomic distance (r = −0.20). In contrast, this relationship was much weaker for negatively co-methylated probe pairs (r = −0.09; Fig. S5c). Age-related changes in pairwise correlation were independent of genomic distance for positively co-methylated probe pairs (r = 0.02), whereas negatively co-methylated probe pairs showed a modest inverse relationship (r = −0.24; Fig. S5d). Co-methylated probe pairs were distributed across all chromosomes, with chromosome 21 containing the fewest co-methylated probe pairs (Fig. S6). In addition, several genomic regions showed several apparent co-methylation hotspots.

Overall, approximately 9% of co-methylated probe pairs (N = 14,928) showed significant age-dependent variability (Bonferroni-adjusted P < 0.05; Fig. S2a), including ~2% that were negatively correlated. Examining top ten most significant ($p<1.0\times{10}^{-43}$) heteroskedastic pairs indicated that covariance between CpGs either remained stable or increased with age, whereas variance at individual CpG sites increased with age (Fig. S2b).

For downstream comparative analyses, the top 1,000 CpG pairs showing the greatest age-related variability ($p<2. 0\times{10}^{-24}$**)** were defined as representative for age-variable co-methylated probe pairs (aVCPs), whereas the bottom 1,000 pairs (i.e., those with the weakest smallest estimated heteroskedasticity; p = ~1.0) were selected to represent age-stable co-methylated probe pairs (aSCPs) (Table S2). Consequently, the aSCPs and aVCPs set comprised 1,375 and 1,432 unique CpGs, respectively. The selected CpGs from both sets were distributed across the genome, although aSCPs appear to form more pronounced local clusters compared to aVCPs (Fig. S4b). The top ranked CpGs within aSCPs showed lower mean methylation levels (~0.2) compared with those within aVCPs (~0.4) (Fig. S7).

Genomic annotation analysis showed that CpGs within both aSCPs and aVCPs were enriched within regulatory domains (Fig. S10). However, aSCPs showed stronger enrichment within promoter regions (OR = 3.1 vs 1.7; P = 8.1 × 10⁻⁸⁹ vs 3.9 × 10⁻²⁰) and CpG islands (OR = 4.6 vs 2.1; P = 1.0 × 10⁻¹⁷¹ vs $p=2.6 \times{10}^{-53}$) and were located near transcription start sites (Table S5).

### Regulatory architecture of age-related co-methylation features

**Overlap with chromatin accessibility**

To evaluate whether age-related variability was constrained through genomic regulation, we overlaid age-related co-methylation CpGs sets with differentially accessible regions (DARs) reported by Patrick, et al. ^1^, who profiled chromatin opening and closing between young and old mice across multiple tissues.

Both aSCPs and aVCPs showed widespread enrichment within both opening and closing DARs across numerous tissues (Fig. S14; Table S11), consistent with chromatin accessibility changes occurring at local regulatory regions. The strongest enrichments for opening DARs were observed in immune cell types, including B cells ($p=2.5 \times{10}^{-89}$), CD4 memory T cells ($p=2.1 \times{10}^{-109}$), and CD8 memory T cells ($p=5.6 \times{10}^{-143}$), whereas enrichment for closing DARs was strongest in adipocytes.

**Histone marks for pairwise and network-level co-methylation**

Distinct histone-modification profiles also emerged when comparing pairwise versus network-level co-methylation features. While network-level age-stable features (aSCMs) were generally enriched for both active and repressive modifications across tissues (Fig. 5), age-stable CpGs identified from pairwise co-methylation (aSCPs) were predominantly enriched for marks associated with active transcription (Fig. S16; Table S14), including H3K27ac, H3K4me3, and H3K9ac. Conversely, aVCPs—mirroring their network-level counterparts (aVCMs)—were consistently enriched for repressive marks (H3K27me3 and H3K9me3) and depleted for active histone modifications, consistent with earlier reports linking these marks to age-associated epigenetic drift ^2,3^.

**Phenotypic relevance of age-related co-methylation features**

We assessed whether genes linked to our age-related co-methylation CpG sets are enriched for genes implicated in GWAS of complex traits, following the approach of Lu, et al. ^4^. For each GWAS trait, genes were ranked according to MAGMA ^5^ gene-level association p-values, and enrichment was evaluated using the top 2.5% of associated genes. When genes linked with the MHC region were excluded from our age-related co-methylation sets in the analysis, we observed varied enrichment of age-related CpG sets across GWAS gene hits (Fig. 7a and Table S16a). For example, age-stable CpG sets (aSCPs and aSCMs) showed over-representation for GWAS gene hits for cardiovascular diseases and aging traits (facial and parental longevity), whereas genes linked to aVCMs were enriched for traits such as the age at menarche, fat and sex hormones.

We also overlapped our age-related CpG sets with EWAS hits for various traits (see methods) and observed notable enrichment patters across age-related co-methylation sets (Fig. 7a and Table S17). Age-stable CpG sets (aSCPs/aSCMs) were consistently enriched for MWAS hits related to autoimmune, inflammatory, developmental, and aging-related phenotypes, associated with early-life origins. In contrast, age-variable features (aVCPs/aVCMs) showed enrichment for traits linked to environmental exposures as well as life-course dynamics, such as smoking, alcohol consumption, and maternal BMI (Table S17).

Moreover, we examined whether genes linked to age-stable and age-variable co-methylation sets overlap within those associated with aging in previous studies. However, we observed no significant enrichment for curated aging-related genes from transcriptomic studies ^6,7^ nor for sex-specific age-associated genes from the GTEx ^8^ dataset, with few exceptions (Fig. S17). These findings are consistent with previous observations that age-related DNA methylation dynamics often do not correspond directly to differential gene expression ^3,9^.

We next tested whether age-stable and aVCMs features overlap with genes responsive to longevity experimental interventions (dietary restriction, feeding timing, fasting, GHR knockout, and Metformin treatment) in mouse and non-human primates ^10-12^. We observed no significant enrichment across all the selected CpG sets from our study (Table S17). This agrees with Haghani, et al. ^13^ observation that methylation modules associated with innate lifespan potential are largely distinct from those responsive to life-extending interventions.

We examined overlap of gernes within co-methylation features with genes linked to epigenetic clocks. We observed marginally significant enrichment (p < 0.05) between genes within aSCMs and those for two second-generation clocks—**DunedinPACE** ^14^ and the Pan-mammalian Clock 3 **(p < 0.05)** ^4^, but no such enrichment for first-generation clocks—primarily designed to capture chronological aging (Fig. 7b and Table S18). Additionally, genes linked to the DunedinPACE clock were also significantly enriched within both age-stable (p < 0.001) and age-variable (p < 0.01) pairwise co-methylated features (aSCPs and aVCPs).

### Sex-stratified co-methylation analysis

We performed sex-stratified network analysis to identify modules with sex-specific preservation trajectories across age groups. We constructed co-methylation modules using the youngest age group from each sex (Age group 1 < 36 years; Fig. S1) and assessed their preservation values across older age groups (Age group 2 to 8) within the same sex, as well as across the opposite sex. Using a reference dataset (Age group 1) from males and females, we identified a total of 83 and 78 modules, respectively (Table S19a,b).

When using male-derived modules as the reference, we observed a moderate decline in cross-sex correlation in module preservation values in early to midlife (i.e., 36 – 55 years), followed by a partial recovery in later age groups (Fig. S18). Notably, the regression lines exhibited increasing ‘tail elongation’ with age, indicating that a subset of modules became progressively more divergent in network preservation between sexes (Fig. S18).

In contrast, when using female-derived modules as reference, cross-sex correlation of preservation statistic remained relatively stable and high across all age groups (r = 0.81–0.90), although the elongation of regression tails with age was still apparent as in male-derived modules (Fig. S18).

#### Sex-biased co-methylation modules based on female reference modules

Several co-methylation modules showed age-related network disintegration within sexes (Fig. S19). To identify modules with pronounced sex differences in age-related co-methylation dynamics, we quantified the absolute change in module preservation statistics between extreme age-groups (i.e., younger and older age groups; Age Group 2 vs. Age Group 8) separately for males and females. Modules were classified as sex-biased if the difference in preservation change between sexes exceeded 5% (Fig. S20). Notably, the correlation between male and female preservation change scores across modules was moderate (r = 0.62; Fig. S21e), indicating that about 38% of the variance in co-methylation remodelling is shared between sexes. These results suggest widespread sex-specific epigenetic restructuring with age consistent with previous work ^15^, which reported that up to 43% of sex-associated CpG sites exhibits age-associated DNA methylation changes.

Using co-methylation modules derived from the female reference dataset, we identified 4 female-biased and 7 male-biased modules with substantial sex differences (>5%; Fig. S20) in preservation statistics between extreme (Age2 vs Age8) age groups (Fig. S21; Table S19a). The larger set of male-biased modules may be partly due to the application of female-derived reference modules to the male dataset. Among the female-biased modules, Module 71 and Module 75 were notable, with their CpGs cognate to *ZNF41/TXNRD2* and *TRIM31/TRIM40* gene clusters, respectively. Region-based functional enrichment showed that Module 71 was enriched (p < 0.001) for biological terms related to development, while Module 75 was enriched for protein phosphorylation (Fig. S21f). These findings are consistent with the involvement of *TRIM* and Zinc family for diverse cellular activities including immune functions and longevity pathways ^16-18^. The top enriched ($p=5.19 \times{10}^{-37}$) gene for Module 71 was *VENTX* that plays a role in regulation of cellular senescence ^19^. Furthermore, this module is associated with neurological disorders in humans, including schizophrenia, bipolar disorder and mood swings (p < 0.001; Table S20a).

Among the male-biased modules, several were characterised by CpGs mapping to gene regions implicated in diverse biological functions (Fig. S21b). Notably, Module 57, exhibited the strongest enrichment for hormonal regulation—particularly insulin signaling ($p=1.0 \times{10}^{-122})$and protein metabolic processes **(**$p=1.14 \times{10}^{-78}$**)** (Fig. S21f and Table S20a), both of which are relevant to aging biology **^20,21^.** Moreover, functional enrichment showed that Module 57 is markedly enriched ($p=1.75 \times{10}^{-145}$) for *NNAT* gene, associated with neurodevelopment and energy metabolism **^22^.** Module 65 was enriched ($p=1.93 \times{10}^{-76}$; Table S19a) for genes involved in arginine methylation—a post-translational modification implicated in chromatin remodelling and transcriptional regulation ^23^. Age-related alterations in arginine methylation have been reported in murine models ^24^, and protein arginine methyltransferases (*PRMTs*) are also considered as potential therapeutic targets in cancer ^25^.

Another striking male-biased module (Module 61) encompassed CpGs mapping to the *HLA* gene cluster, a region involved in immune regulation, genetic imprinting, and embryonic development (Fig. S21f). Finally, Module 53, also male-biased, featured CpGs proximal to *GLI2* gene, a key transcription factor in the Hedgehog signaling pathway linked to neurodevelopment and stem cell maintenance **^26^**, consistent with our findings (Table S20a). Overall, the pronounced age-related changes in these modules in males than females suggest sex-specific trajectories for immune, metabolic, and developmental regulation with aging, potentially contributing to observed epidemiological differences in aging phenotypes between men and women.

#### Sex-biased co-methylation modules based on male reference modules

Using co-methylation modules derived from the male reference dataset, we observed a lower correlation between male and female preservation change scores across modules (r = 0.55, compared to r = 0.62 when using the female reference), implying a greater degree of sex-specific network reorganization when male-derived epigenetic architecture is used as the reference. This is consistent with the previous work showing that larger number of CpG sites in males show age-related increases in methylation variability than females ^15^. We identified 5 male-biased (|Δ preservation statistics| > 5%; Fig. S20a) and 8 female-biased modules (Fig. S21b and Table S19b). Notably, three male-biased modules—Module 52 (enriched for developmental processes), Module 59 (linked to hormonal regulation and metabolism), and Module 62 (associated with immune system function)—were consistently classified as male-biased in both male- and female-derived reference modules highlighting a consistent sex-specific epigenetic remodelling with age.

Module 64, another female-biased module, included members of the zinc finger protein family, a finding consistent with its female bias in the female-derived reference network discussed above. Module 77 stands out for its CpGs mapping to the *CBX5/SUMG1* gene cluster. *CBX5* (also known as *HP1α*) is a key regulator of heterochromatin organization, cellular senescence, and lifespan regulation ^27^. Partial loss of *CBX5* in drosophila reduces lifespan by over 50% ^28^. Module 74 included CpGs near the *GADD45B/GNG7* gene cluster. Among these, *GADD45B* is of particular interest given its role in DNA demethylation, cellular stress response, and senescence ^29,30^.

Module 80, which includes CpGs mapping to the *PTPRN2/DNAJB6* gene cluster, showed strong enrichment for pathways related to placental development (Fig. S21c) consistent with the hypothesis of female-specific remodelling in reproductive or developmentally imprinted loci. Notably, *PTPRN2* has been associated with puberty timing and age at menarche in GWAS of Japanese populations ^31^ and shows sex-specific methylation patterns in Parkinson’s disease, with hypomethylation in female brains and hypermethylation in male brains ^32^.

Interestingly, we observed the lowest concordance in module preservation statistics between males and females around midlife (50–55 years) when using male-derived reference modules (Fig. S18; r = 0.66). This drop in correlation suggests a period of increased sex divergence in co-methylation architecture, likely driven by the impact of midlife hormonal transitions, such as menopause in females and gradual androgen decline in males, both of which have been associated with epigenetic remodelling with aging ^33,34^. As such, to identify sex-specific modules potentially associated with these changes, we calculated the absolute change in preservation statistics between the younger age group (age group 2; 36–44 years) and midlife (age group 4; 50–55 years; Fig. 1). Consequently, we identified four male-biased and two female-biased modules showing a substantial (>5%; Fig. S20c) difference in module preservation change in one sex relative to the other (Fig. S22a and Table S21).

Male-biased Module 59 included CpGs within *NNAT* gene ($p=4.2 \times{10}^{-147}$; Table S21) (discussed earlier)—a paternally expressed imprinted gene linked to neurodevelopment and disorders including schizophrenia ^35^. This module was also strongly enriched for hormonal regulation and energy metabolism axis (Fig. S22c and Table S22). In contrast, the striking female-biased Module 43 was characterised by CpGs within the *GNAS* gene)—a maternally expressed imprinted locus involved in hormone signaling and developmental pathways (Fig. S22c). Additionally, female-biased Module 63 included CpGs within *HOX* genes, which are well known for driving developmental processes.

### References

1	Patrick, R. *et al.* The activity of early-life gene regulatory elements is hijacked in aging through pervasive AP-1-linked chromatin opening. *Cell Metab* **36**, 1858-1881.e1823 (2024). <https://doi.org/10.1016/j.cmet.2024.06.006>

2	Slieker, R. C. *et al.* Age-related accrual of methylomic variability is linked to fundamental ageing mechanisms. *Genome Biol* **17**, 191 (2016). <https://doi.org/10.1186/s13059-016-1053-6>

3	Reynolds, L. M. *et al.* Age-related variations in the methylome associated with gene expression in human monocytes and T cells. *Nat Commun* **5**, 5366 (2014). <https://doi.org/10.1038/ncomms6366>

4	Lu, A. T. *et al.* Universal DNA methylation age across mammalian tissues. *Nat Aging* **3**, 1144-1166 (2023). <https://doi.org/10.1038/s43587-023-00462-6>

5	de Leeuw, C. A., Mooij, J. M., Heskes, T. & Posthuma, D. MAGMA: generalized gene-set analysis of GWAS data. *PLoS Comput Biol* **11**, e1004219 (2015). <https://doi.org/10.1371/journal.pcbi.1004219>

6	de Magalhães, J. P. *et al.* Human Ageing Genomic Resources: updates on key databases in ageing research. *Nucleic acids research* **52**, D900-D908 (2024).

7	Aging Atlas Consortium. Aging Atlas: a multi-omics database for aging biology. *Nucleic acids research* **49**, D825-D830 (2021).

8	Wang, S., Dong, D., Li, X. & Wang, Z. Pan-tissue Transcriptome Analysis Reveals Sex-dimorphic Human Aging. *bioRxiv*, 2023.2005. 2026.542373 (2023).

9	Horvath, S. DNA methylation age of human tissues and cell types. *Genome Biol* **14**, R115 (2013). <https://doi.org/10.1186/gb-2013-14-10-r115>

10	Acosta-Rodríguez, V. *et al.* Circadian alignment of early onset caloric restriction promotes longevity in male C57BL/6J mice. *Science* **376**, 1192-1202 (2022).

11	Sun, L. Y. *et al.* Growth hormone-releasing hormone disruption extends lifespan and regulates response to caloric restriction in mice. *Elife* **2**, e01098 (2013).

12	Yang, Y. *et al.* Metformin decelerates aging clock in male monkeys. *Cell* **187**, 6358-6378. e6329 (2024).

13	Haghani, A. *et al.* DNA methylation networks underlying mammalian traits. *Science* **381**, eabq5693 (2023). <https://doi.org/10.1126/science.abq5693>

14	Belsky, D. W. *et al.* DunedinPACE, a DNA methylation biomarker of the pace of aging. *Elife* **11**, e73420 (2022).

15	Yusipov, I. *et al.* Age-related DNA methylation changes are sex-specific: a comprehensive assessment. *Aging (Albany NY)* **12**, 24057-24080 (2020). <https://doi.org/10.18632/aging.202251>

16	Ozato, K., Shin, D. M., Chang, T. H. & Morse, H. C., 3rd. TRIM family proteins and their emerging roles in innate immunity. *Nat Rev Immunol* **8**, 849-860 (2008). <https://doi.org/10.1038/nri2413>

17	Hamilton, B. *et al.* A systematic RNAi screen for longevity genes in C. elegans. *Genes Dev* **19**, 1544-1555 (2005). <https://doi.org/10.1101/gad.1308205>

18	Cassandri, M. *et al.* Zinc-finger proteins in health and disease. *Cell Death Discov* **3**, 17071 (2017). <https://doi.org/10.1038/cddiscovery.2017.71>

19	Wu, X. *et al.* VentX trans-activates p53 and p16ink4a to regulate cellular senescence. *J Biol Chem* **286**, 12693-12701 (2011). <https://doi.org/10.1074/jbc.M110.206078>

20	Tatar, M., Bartke, A. & Antebi, A. The endocrine regulation of aging by insulin-like signals. *Science* **299**, 1346-1351 (2003). <https://doi.org/10.1126/science.1081447>

21	Soultoukis, G. A. & Partridge, L. Dietary Protein, Metabolism, and Aging. *Annu Rev Biochem* **85**, 5-34 (2016). <https://doi.org/10.1146/annurev-biochem-060815-014422>

22	Joseph, R. M. Neuronatin gene: Imprinted and misfolded: Studies in Lafora disease, diabetes and cancer may implicate NNAT-aggregates as a common downstream participant in neuronal loss. *Genomics* **103**, 183-188 (2014). <https://doi.org/10.1016/j.ygeno.2013.12.001>

23	Blanc, R. S. & Richard, S. Arginine Methylation: The Coming of Age. *Mol Cell* **65**, 8-24 (2017). <https://doi.org/10.1016/j.molcel.2016.11.003>

24	Zhang, F. *et al.* Global analysis of protein arginine methylation. *Cell Rep Methods* **1**, 100016 (2021). <https://doi.org/10.1016/j.crmeth.2021.100016>

25	Hwang, J. W., Cho, Y., Bae, G. U., Kim, S. N. & Kim, Y. K. Protein arginine methyltransferases: promising targets for cancer therapy. *Exp Mol Med* **53**, 788-808 (2021). <https://doi.org/10.1038/s12276-021-00613-y>

26	Ingham, P. W. & McMahon, A. P. Hedgehog signaling in animal development: paradigms and principles. *Genes Dev* **15**, 3059-3087 (2001). <https://doi.org/10.1101/gad.938601>

27	Mendelsohn, A. R. & Larrick, J. W. Stem Cell Depletion by Global Disorganization of the H3K9me3 Epigenetic Marker in Aging. *Rejuvenation Res* **18**, 371-375 (2015). <https://doi.org/10.1089/rej.2015.1742>

28	Larson, K. *et al.* Heterochromatin formation promotes longevity and represses ribosomal RNA synthesis. *PLoS Genet* **8**, e1002473 (2012). <https://doi.org/10.1371/journal.pgen.1002473>

29	Moskalev, A. A. *et al.* Gadd45 proteins: relevance to aging, longevity and age-related pathologies. *Ageing Res Rev* **11**, 51-66 (2012). <https://doi.org/10.1016/j.arr.2011.09.003>

30	Magimaidas, A. *et al.* Gadd45b deficiency promotes premature senescence and skin aging. *Oncotarget* **7**, 26935-26948 (2016). <https://doi.org/10.18632/oncotarget.8854>

31	Horikoshi, M. *et al.* Elucidating the genetic architecture of reproductive ageing in the Japanese population. *Nat Commun* **9**, 1977 (2018). <https://doi.org/10.1038/s41467-018-04398-z>

32	Kochmanski, J., Kuhn, N. C. & Bernstein, A. I. Parkinson's disease-associated, sex-specific changes in DNA methylation at PARK7 (DJ-1), SLC17A6 (VGLUT2), PTPRN2 (IA-2β), and NR4A2 (NURR1) in cortical neurons. *NPJ Parkinsons Dis* **8**, 120 (2022). <https://doi.org/10.1038/s41531-022-00355-2>

33	Levine, M. E. *et al.* Menopause accelerates biological aging. *Proc Natl Acad Sci U S A* **113**, 9327-9332 (2016). <https://doi.org/10.1073/pnas.1604558113>

34	Wilkinson, H. N. & Hardman, M. J. The role of estrogen in cutaneous ageing and repair. *Maturitas* **103**, 60-64 (2017). <https://doi.org/10.1016/j.maturitas.2017.06.026>

35	Tesfaye, M. *et al.* Sex effects on DNA methylation affect discovery in epigenome-wide association study of schizophrenia. *Mol Psychiatry* **29**, 2467-2477 (2024). <https://doi.org/10.1038/s41380-024-02513-9>
