## Supplementary Figures for "Age-related dynamics of DNA co-methylation modules in humans"

Supplementary Table S1. Glossary of abbreviations used in the manuscript

| **Abbreviation** | **Definition** |
| --- | --- |
| DNAm | DNA methylation |
| CpG | Cytosine–phosphate–guanine dinucleotide |
| aDMP | Age-associated differentially methylated position |
| aVMP | Age-associated variably methylated position |
| aSCPs | Age-stable co-methylated probe pairs |
| aVCPs | Age-variable co-methylated probe pairs |
| aSCMs | Age-stable co-methylation modules |
| aVCMs | Age-variable co-methylation modules |
| WGCNA | Weighted Gene Co-expression Network Analysis |
| GAM | Generalized Additive Model |
| meQTL | Methylation quantitative trait locus |
| cis-meQTL | Cis methylation quantitative trait locus |
| trans-meQTL | Trans methylation quantitative trait locus |
| PPI | Protein–protein interaction |
| TFBS | Transcription factor binding site |
| TSS | Transcription start site |
| TAD | Topologically associating domain |
| Hi-C | High-throughput chromosome conformation capture |
| DAR | Differentially accessible region |
| PMD | Partially methylated domain |
| HMD | Highly methylated domain |
| GWAS | Genome-wide association study |
| EWAS | Epigenome-wide association study |
| MAGMA | Multi-marker Analysis of GenoMic Annotation |
| GREAT | Genomic Regions Enrichment of Annotations Tool |
| STRING | Search Tool for the Retrieval of Interacting Genes/Proteins |
| GoDMC | Genetics of DNA Methylation Consortium |
| ENCODE | Encyclopedia of DNA Elements |
| OREML | Omics-based Restricted Maximum Likelihood |
| OSCA | Omic-data-based Complex Trait Analysis |
| OR | Odds ratio |
| CI | Confidence interval |
| TF | Transcription factor |
| CBX8 | Chromobox protein homolog 8 |
| DR | Dietary restriction |
| GHR | Growth hormone receptor |

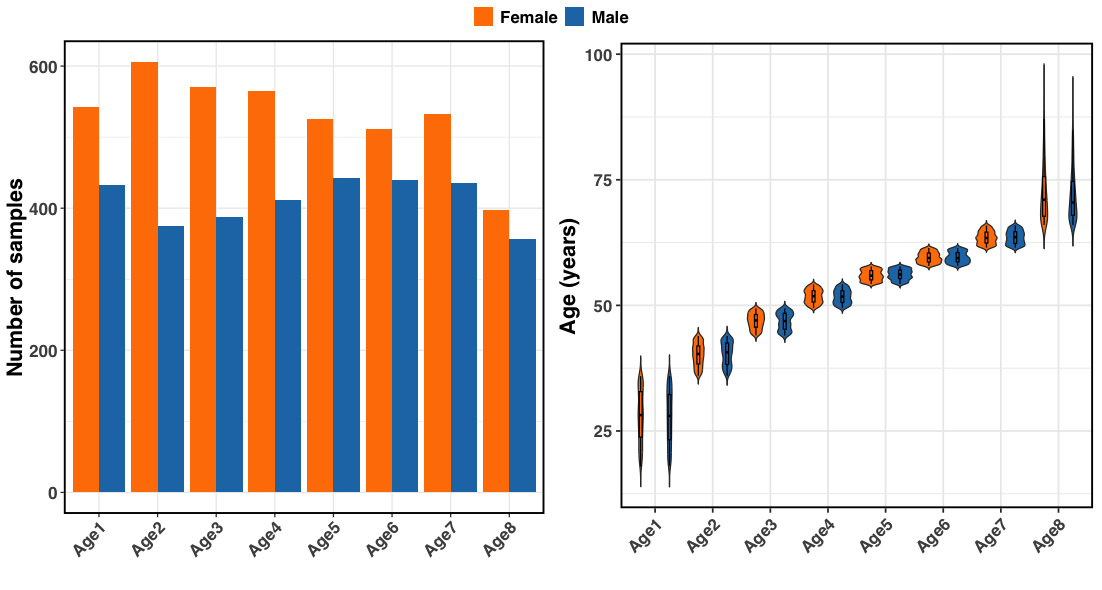

Supplementary Figure 1: Age distribution for males and females across eight defined age groups. While the proportion of males was slightly lower than females in all age groups, the overall age distribution was comparable between sexes.

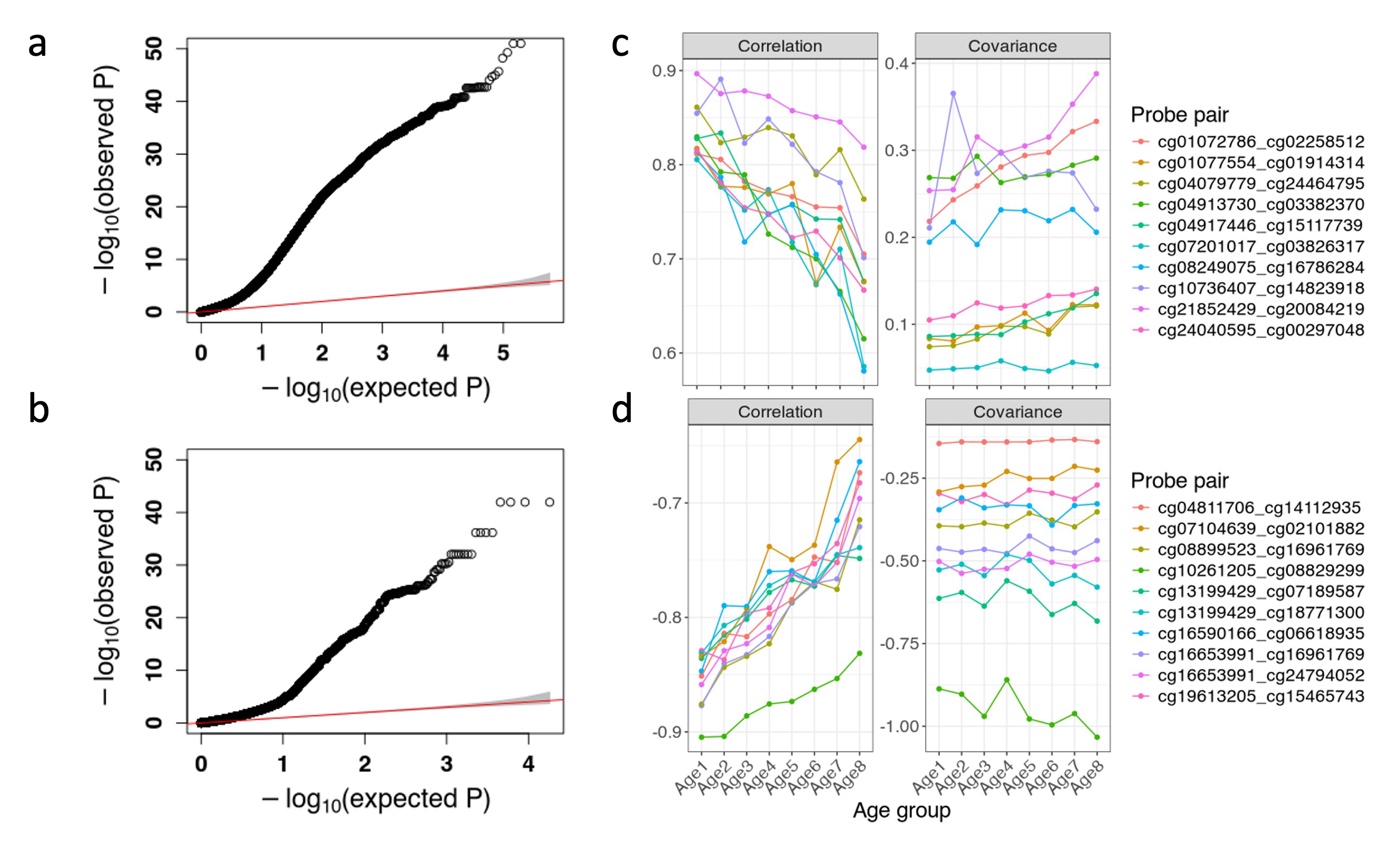

Supplementary Figure 2: Discovery of co-methylated probe pairs and correlation and covariance dynamics.

**(a, b)** Q–Q plots of p-values from heteroskedasticity tests assessing age-related changes in pairwise correlation strength for positively (top) and negatively correlated probe pairs (bottom).

**(c)** Positively correlated probe pairs; **(d)** negatively correlated probe pairs. While correlation strength tends to decline with age, covariance either increases or remains stable. The top 10 age-variable probes were ranked based on heteroskedasticity test significance.

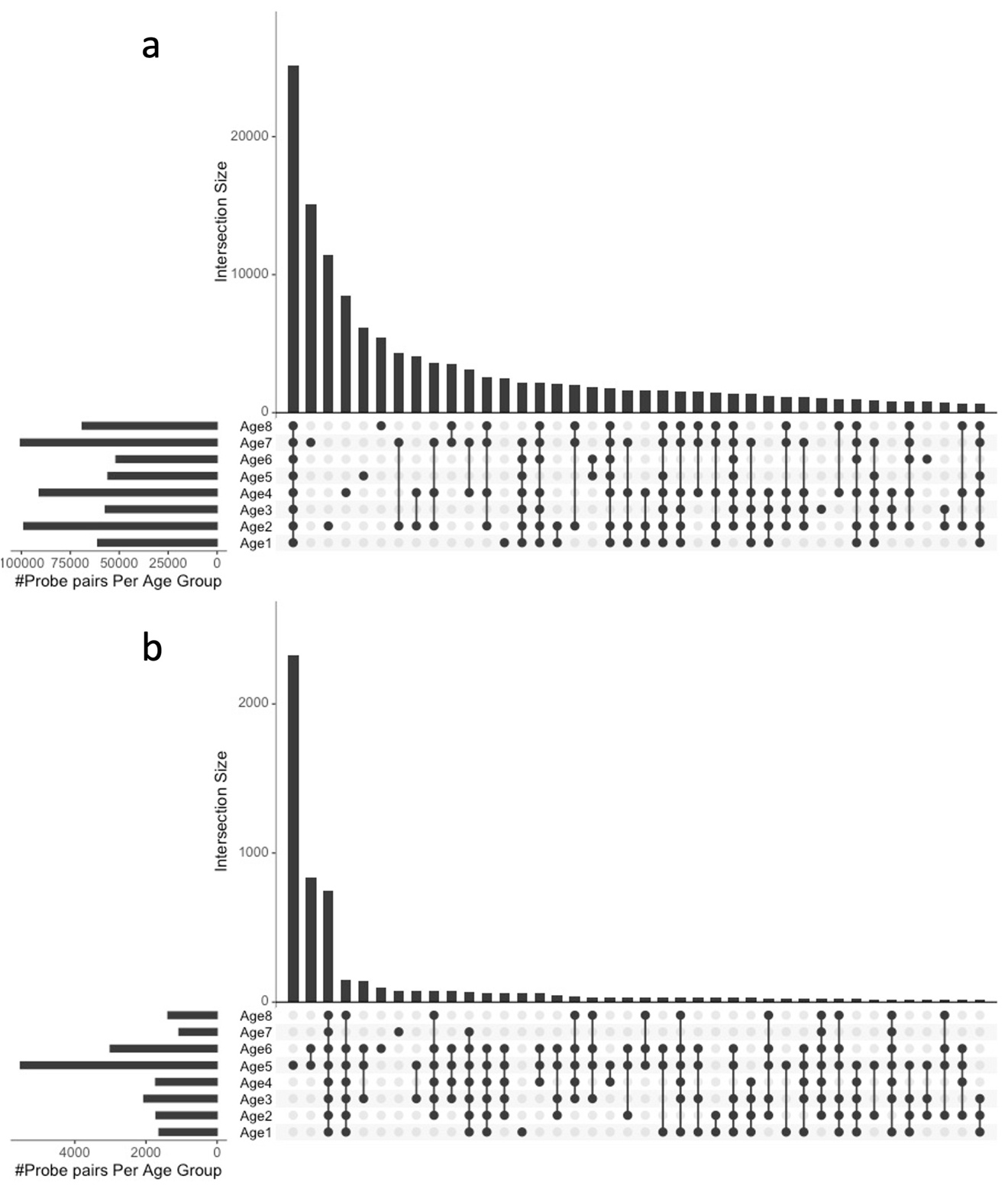

Supplementary Figure 3: Overlaps of correlated probe pairs.

Overlap of positively (a; r < -0.80) and negatively (b; r > 0.80) correlated CpG probe pairs across study age groups (Age1–Age8), based on pairwise correlations computed within each group. The bar on the left of the plot shows the number of correlated probe pairs for each age group and the top bars shows the overlap size.

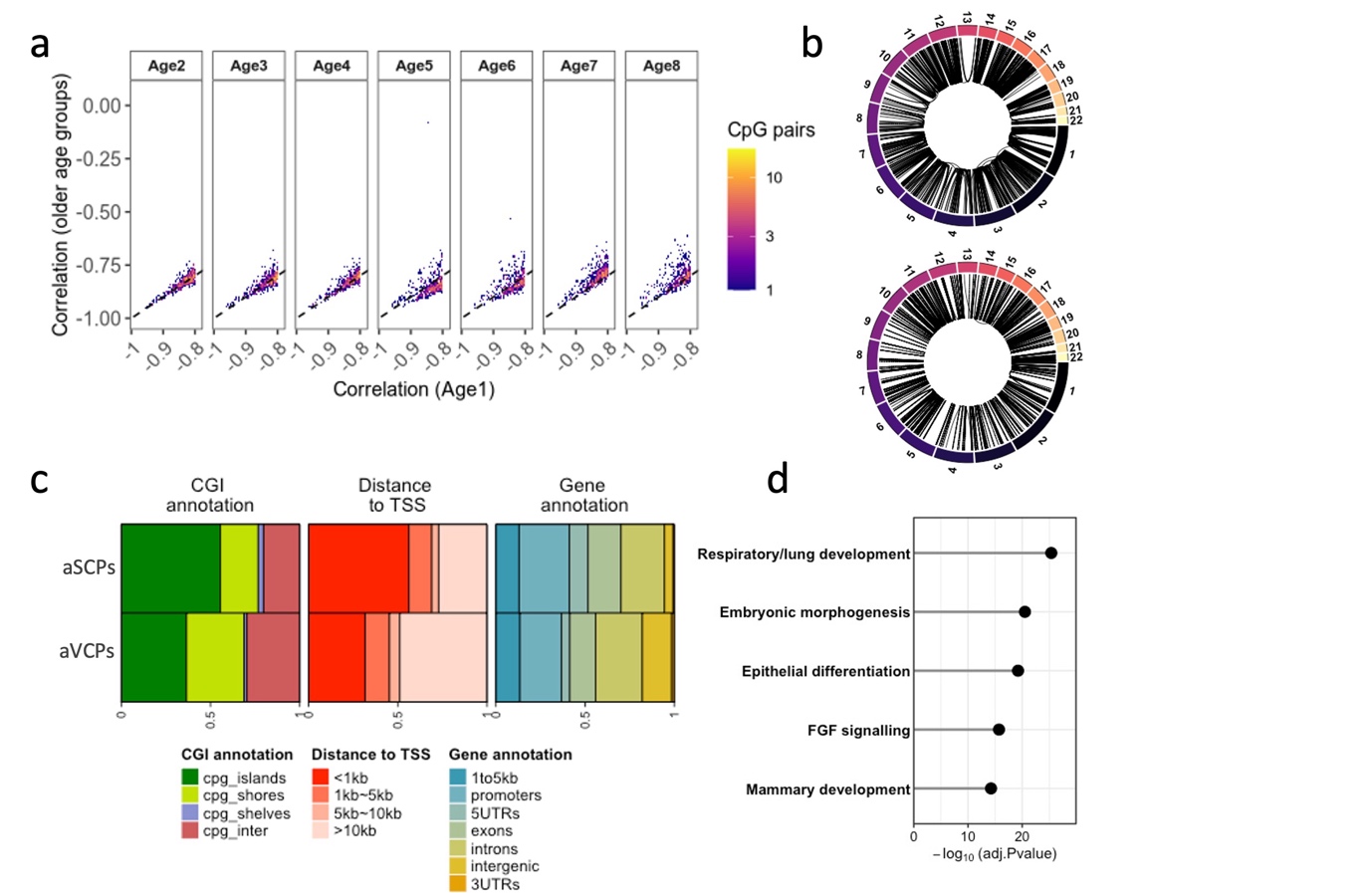

Supplementary Figure 4: Discovery of age-stable and age-variable co-methylated probe pairs (aSCPs and aVCPs) based on heteroskedasticity test.

**(a)** Density heatmaps of negatively correlated CpG probe pairs (r < -0.80) identified in the youngest age group (Age1) and tracked across older age groups. Each panel compares correlation values in Age1 with those observed in subsequent age groups. Colour intensity represents the density of CpG probe pairs within each bin.

**(b)** The distribution of CpGs within the aSCPs (top) and aVCPs (bottom) co-methylation sets across the genome.

**(c)** Genomic annotations of aSCPs and aVCPs CpG sets. The genomic categories are characterised into CpG (CGI), distance to the transcription start sites (TSS) and genic annotations.

**(d)** Major functional themes associated with aVCPs. No significant (p < 0.05) biological themes were linked to aSCPs. The detailed biological pathways are provided in Table S14b.

Abbreviations: age-stable co-methylated probe pairs (aSCPs), age-variable co-methylated probe pairs (aVCPs).

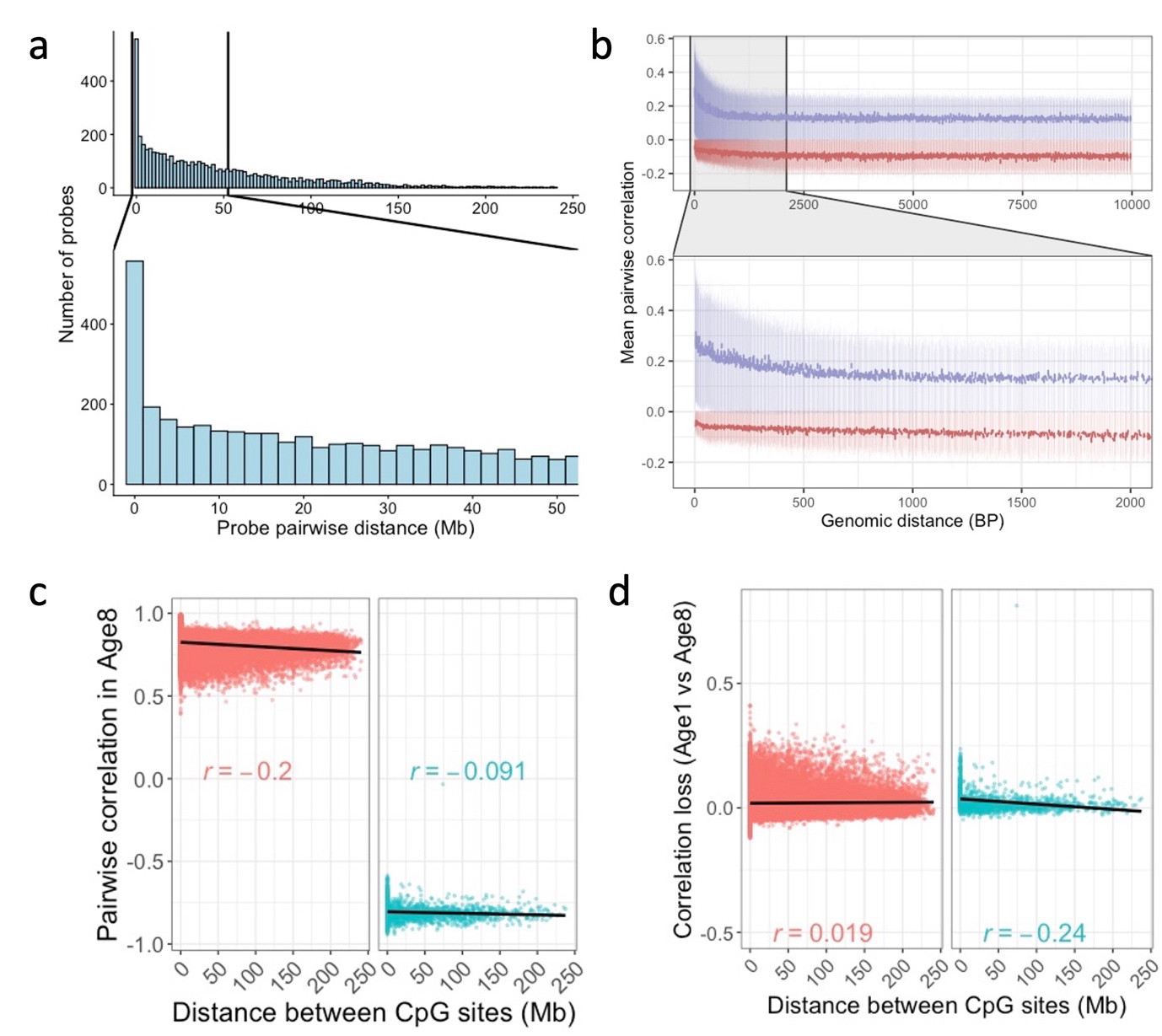

Supplementary Figure 5: Genomic distance profile for co-methylated CpG probe pairs.

**(a)** Histogram of pairwise distance across all positively and negatively correlated probes pairs. We observed that a large proportion of co-methylated probe-pairs are less than < 1Mb but some co-methylated pairs are > 200Mb.

**(b)** Decay of pairwise correlations for positively (r > 0.80; blue) and negatively (r < –0.80; red) correlated probe pairs with increasing genomic distance. Co-methylated probe pairs ≤10 kb apart were binned into 200 intervals based on distance, and mean correlation ± SD was computed per bin. A rapid decline in co-methylation is observed around 1 kb, beyond which correlation levels plateau. Inset shows a zoomed view of the 0–2 kb region.
**(c)** Relationship between genomic distance and pairwise correlation in the oldest age group (Age8) for CpG pairs that were positively or negatively correlated in the youngest age group (Age1).
**(e)** Relationship between genomic distance and the change in pairwise correlation magnitude from the youngest (Age1) to the oldest (Age8) age group for positive and negative co-methylated CpG pairs.

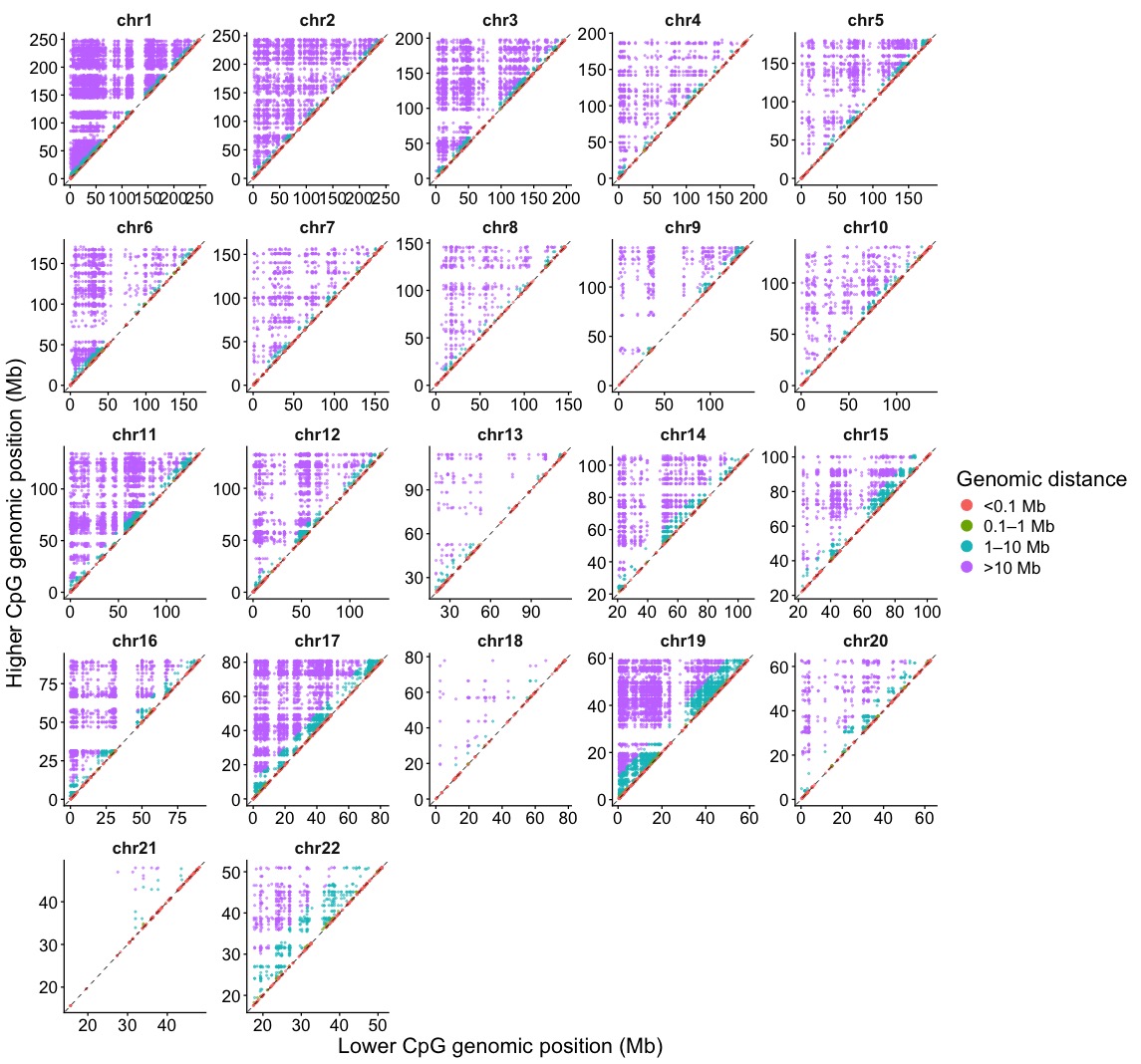

Supplementary Figure 6: Spatial distribution of co-methylated CpG pairs across individual chromosomes. CpG pairs were plotted according to the genomic coordinates of the lower and higher positioned CpG within each chromosome and coloured by genomic separation.

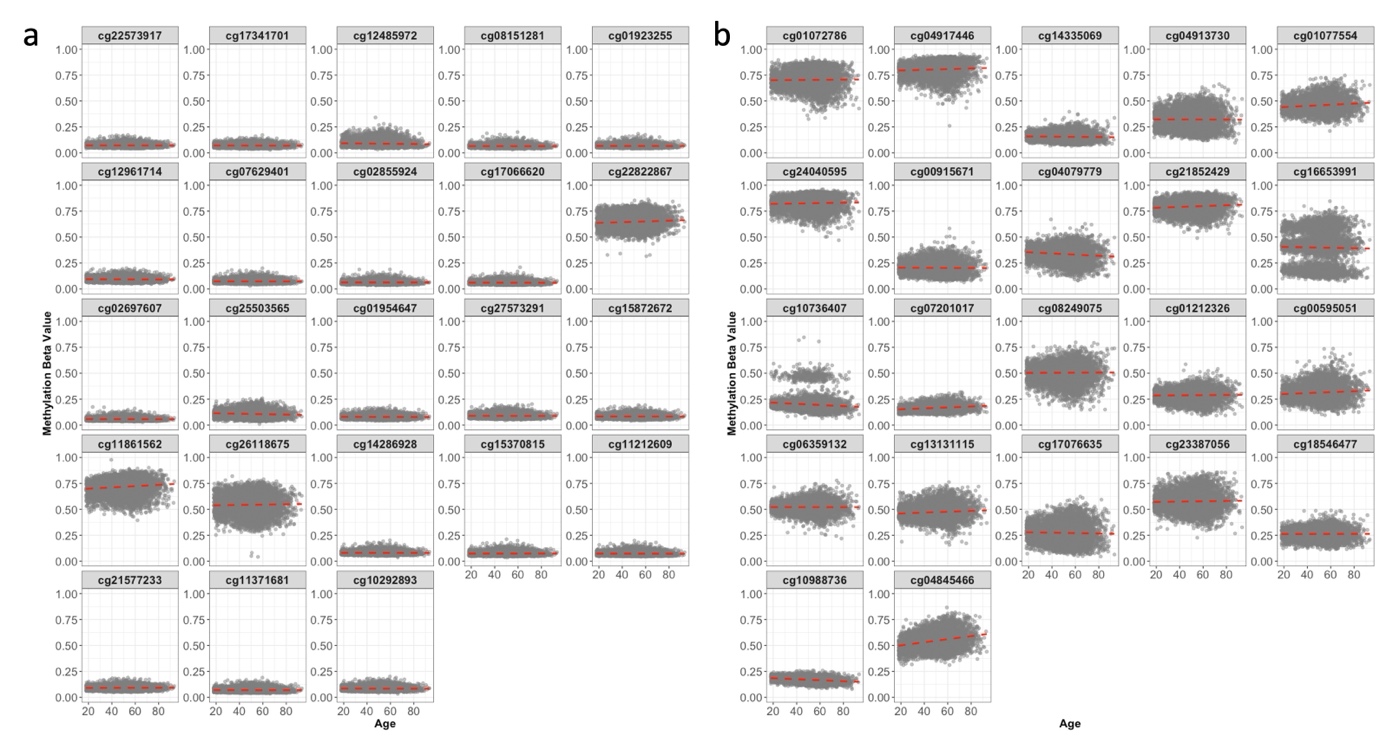

Supplementary Figure 7: DNA methylation profiles of selected top CpGs (ranked by heteroskedasticity test significance p-value) within aSCPs and aVCPs sets.
**(a)** Age trajectories for the top age-stable co-methylated probe pairs (aSCPs)

**(b)** Top age-variable co-methylated probe pairs (aVCPs). aSCPs generally showed lower overall methylation levels and maintained consistent patterns across age groups. In contrast, aVCPs displayed increased methylation variability with age in line with age-associated epigenetic drift. One CpG within the FGFR2/WDR11-associated pair (cg16653991) is a previously reported cis-mQTL target ^1^.

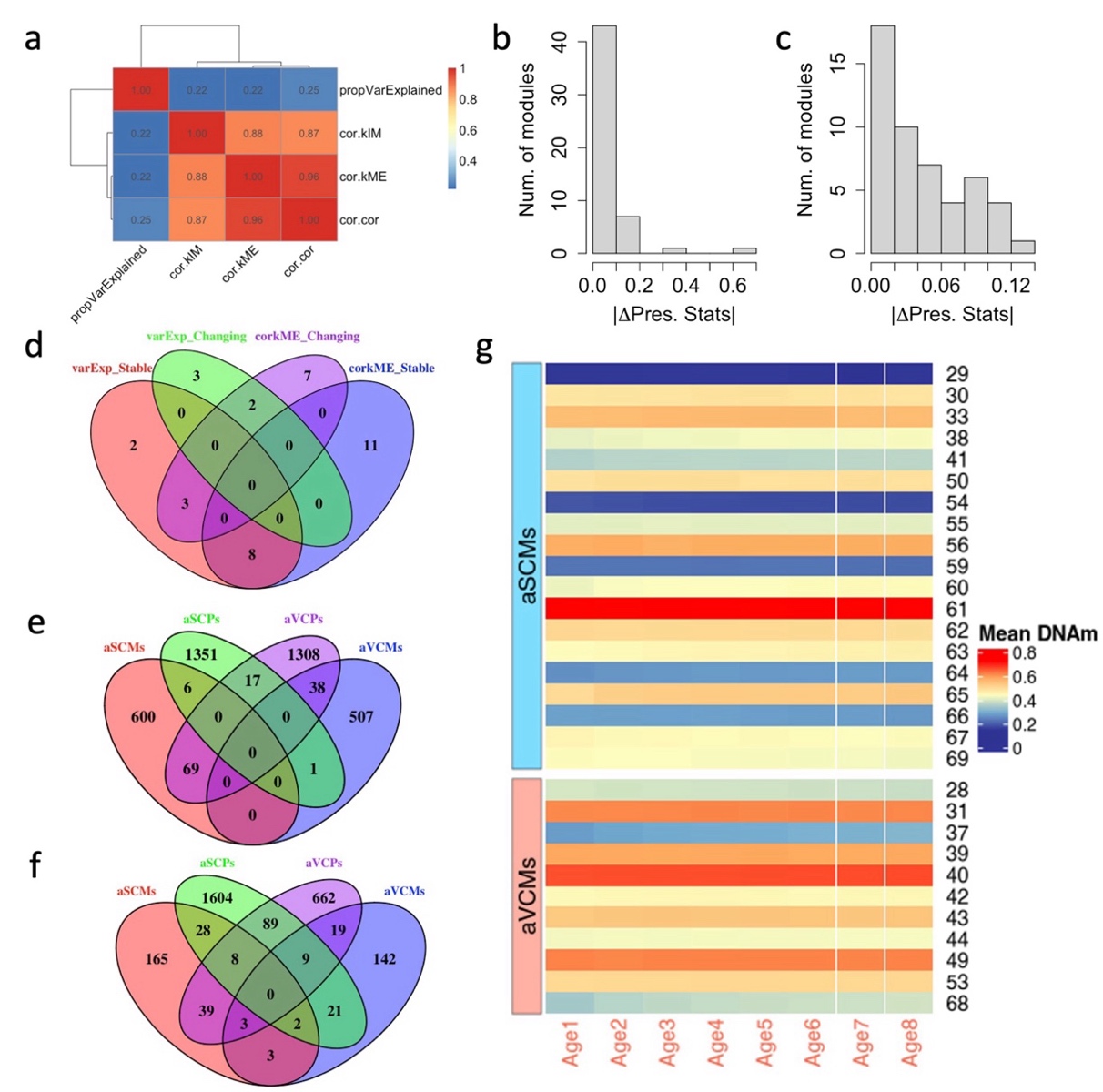

Supplementary Figure 8: Correlations among WGCNA preservation statistics and overlaps of selected set of CpGs and genes.
(a) Pairwise correlations among WGCNA module preservation statistics across age groups. Strong correlations were observed among network connectivity-based metrics—cor.kME, cor.cor, and cor.kIM (r > 0.85)—while the proportion of variance explained (propVarExp) showed weak correlations with these measures (r < 0.25). These results suggest that connectivity and variance metrics capture distinct aspects of module preservation. Therefore, cor.kME and propVarExp were selected as complementary metrics for downstream analyses.

**(b, c)** Distribution of changes in preservation statistics between extreme age groups (Age group 2 vs. Age group 8) for cor.kME (b) and propVarExp (c).

**(d)** Overlap of candidate age-stable and age-variable modules identified using cor.kME and propVarExp. Modules showing discordant classification across metrics (n = 3) were not assigned to either category and were excluded from downstream analyses.
**(e)** Overlap between CpG sites identified using heteroskedasticity-based (aSCPs and aVCPs) and WGCNA-based (aSCMs and aVCMs) approaches. Minimal overlap was observed, suggesting that these methods capture distinct dimensions of epigenetic variation—local (pairwise) versus global (network-level) co-methylation structure.
**(f)** Overlap of genes associated with age-stable and age-variable CpG sites across both heteroskedasticity and WGCNA analyses. While overlap at the CpG level was limited, a notable number of genes were shared between age-stable and age-variable CpG sets (e.g., N = 153 between aSCPs vs aVCPs). This suggests the coexistence of conserved and dynamic methylation patterns within the same gene loci.

**(g)** Mean methylation levels of CpGs within age-stable and age-variable modules from WGCNA. The number represent module identifiers within aSCPs and aVCPs module sets.

Abbreviations: age-stable co-methylation modules (aSCMs), age-variable co-methylation modules (aVCMs), age-stable co-methylated probe pairs (aSCPs), age-variable co-methylated probe pairs (aVCPs).

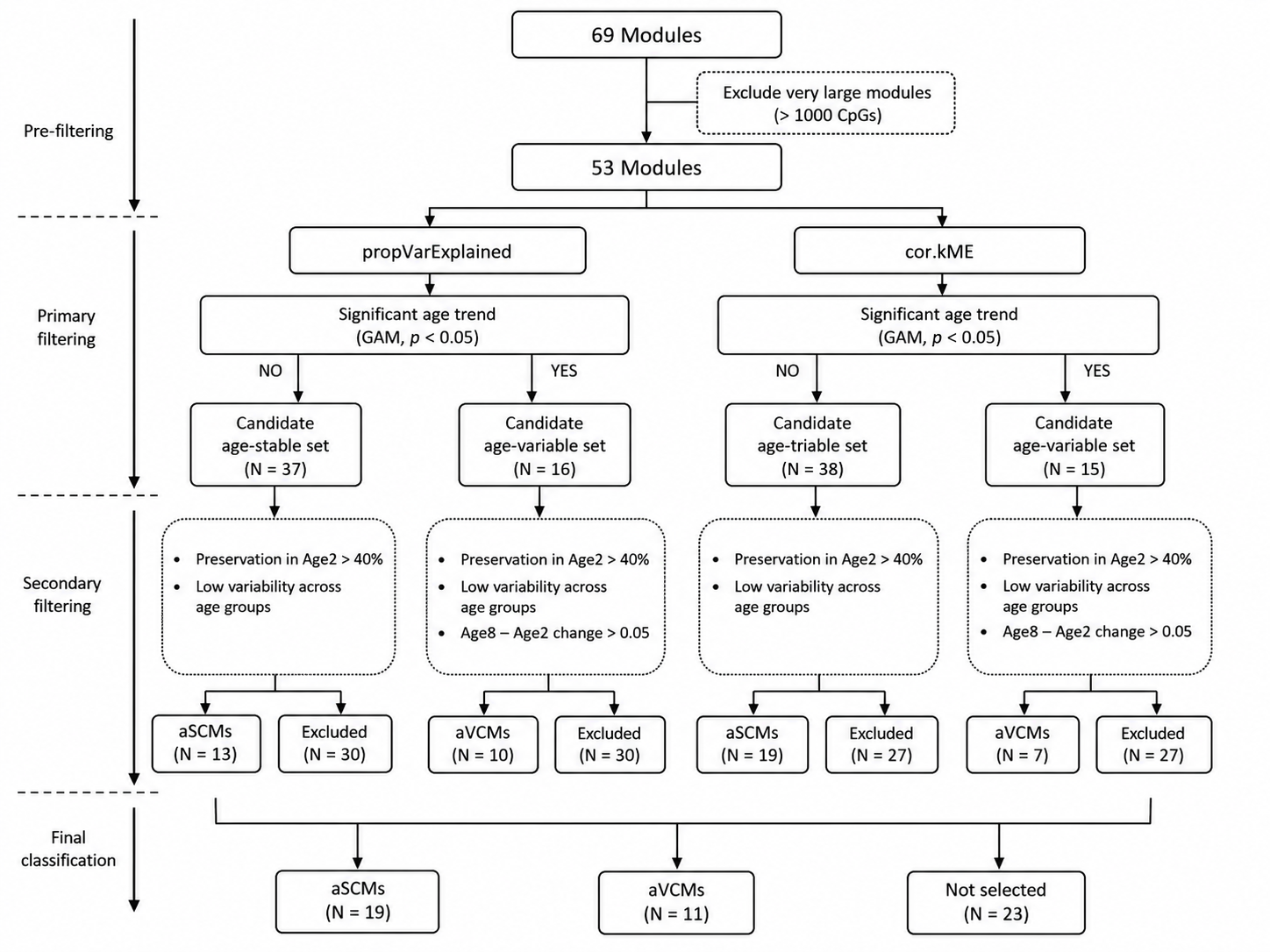

Supplementary Figure 9: Overview of module selection.

DNA co-methylation networks identified using WGCNA ^2^ were first filtered to remove very large modules (>1,000 CpGs), leaving 53 modules for downstream analyses. For each preservation metric (propVarExplained and cor.kME), generalized additive models (GAMs) were fitted across age groups to identify candidate age-stable and age-variable modules based on a significant age trend (p < 0.01). Candidate age-stable modules were required to show strong preservation in the youngest test age group (Age2; preservation > 0.40) and low variability across age groups. Candidate age-variable modules were required to meet the same preservation criterion in Age2 and exhibit a preservation change between Age2 and Age8 greater than 0.05. Modules satisfying these criteria were classified as age-stable (aSCMs) or age-variable (aVCMs) for each preservation metric. Finally, modules showing consistent classification across either preservation metric were retained, whereas inconsistently classified modules and module-size outliers were excluded, resulting in 19 aSCMs and 11 aVCMs. Abbreviations: age-stable co-methylation modules (aSCMs), age-variable co-methylation modules (aVCMs), age-stable co-methylated probe pairs (aSCPs), age-variable co-methylated probe pairs (aVCPs).

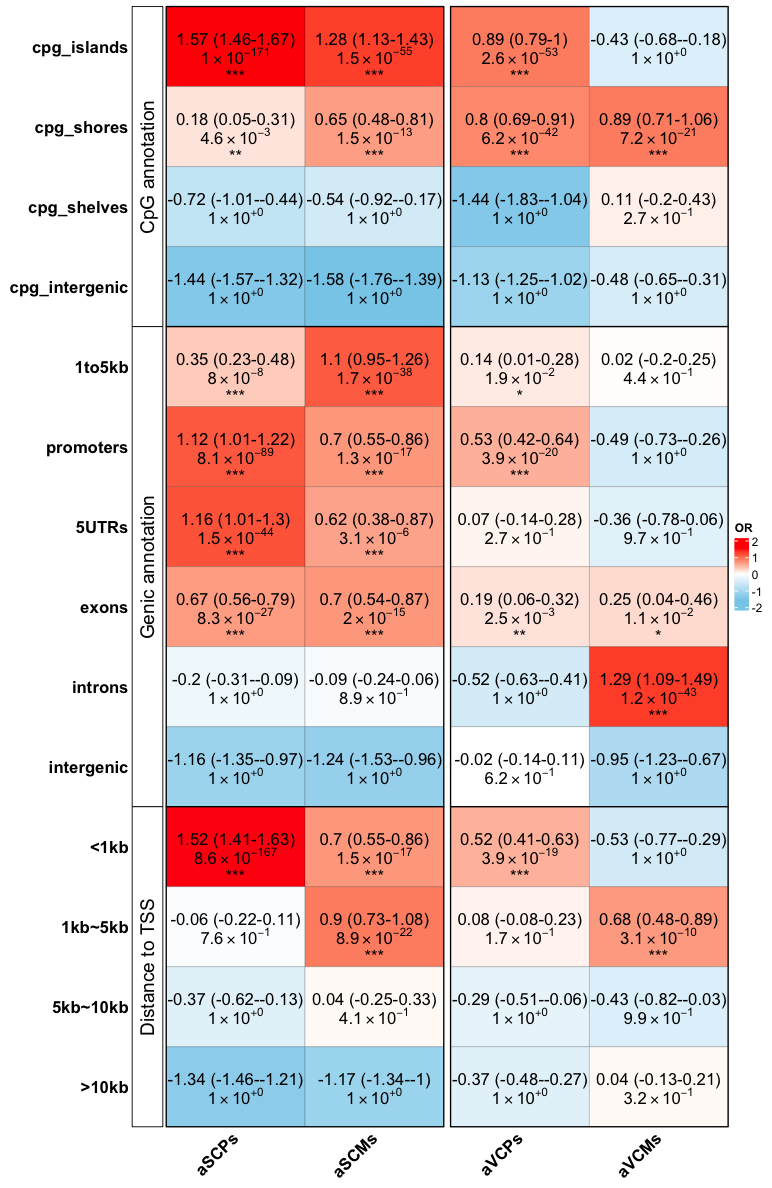

Supplementary Figure 10: Enrichment of age-related co-methylation sets across different genomic domains. The bars are coloured based on log(OR). OR < 0 indicate depletion, while OR > 0, represent over-representation. The asterisks are the significance strength (* = p < 0.05; ** p < 0.01; *** = p < 0.001). The detailed results are provided in Table S4.

Abbreviations: age-stable co-methylation modules (aSCMs), age-variable co-methylation modules (aVCMs), age-stable co-methylated probe pairs (aSCPs), age-variable co-methylated probe pairs (aVCPs).

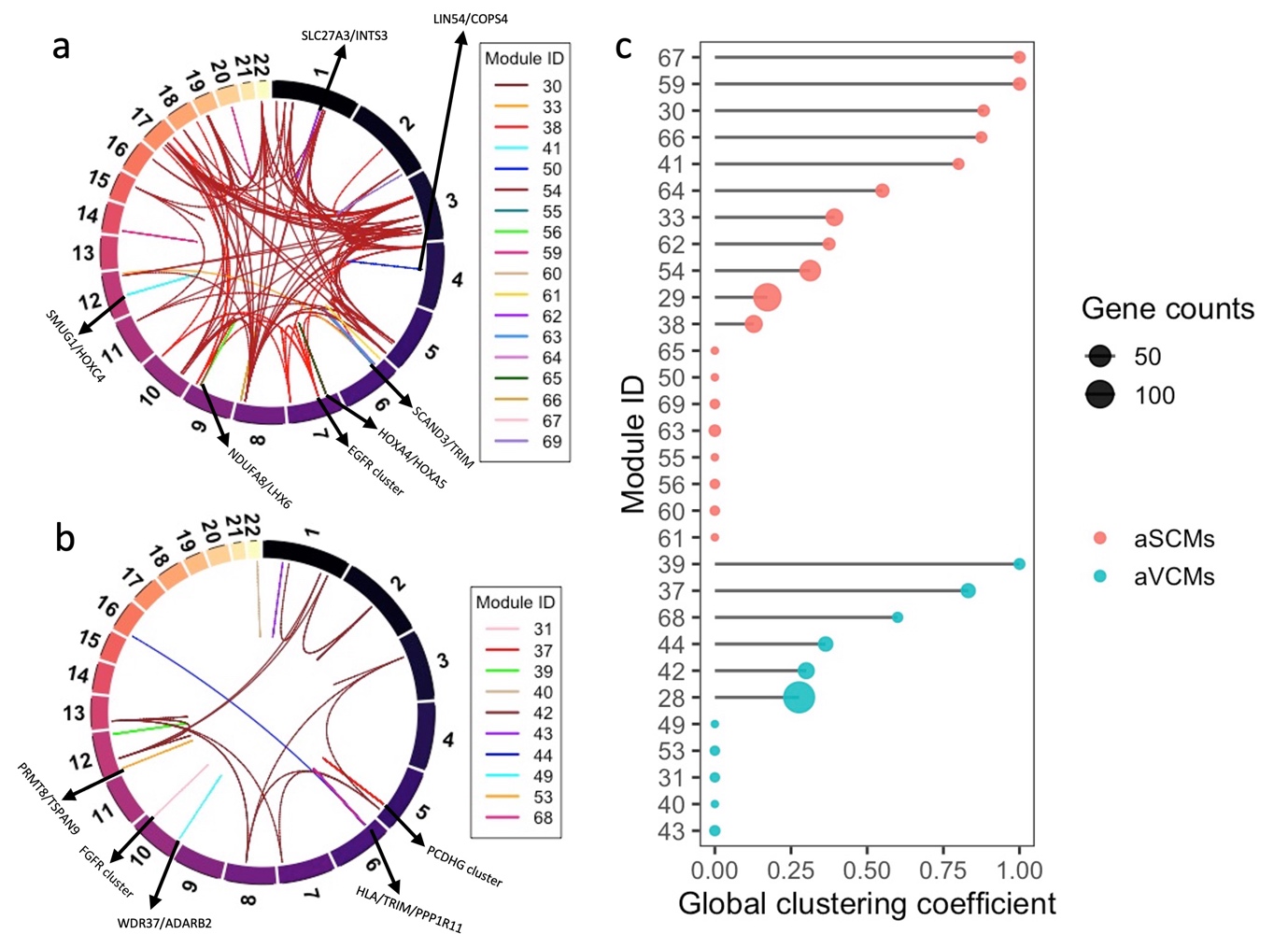

Supplementary Figure 11: Protein–protein interaction (PPI) networks derived from WGCNA for age-stable (aSCMs; **a**) and age-variable (aVCMs; **b**) modules; (**c**) Global clustering coefficient estimates across modules.

Abbreviations: age-stable co-methylation modules (aSCMs), age-variable co-methylation modules (aVCMs).

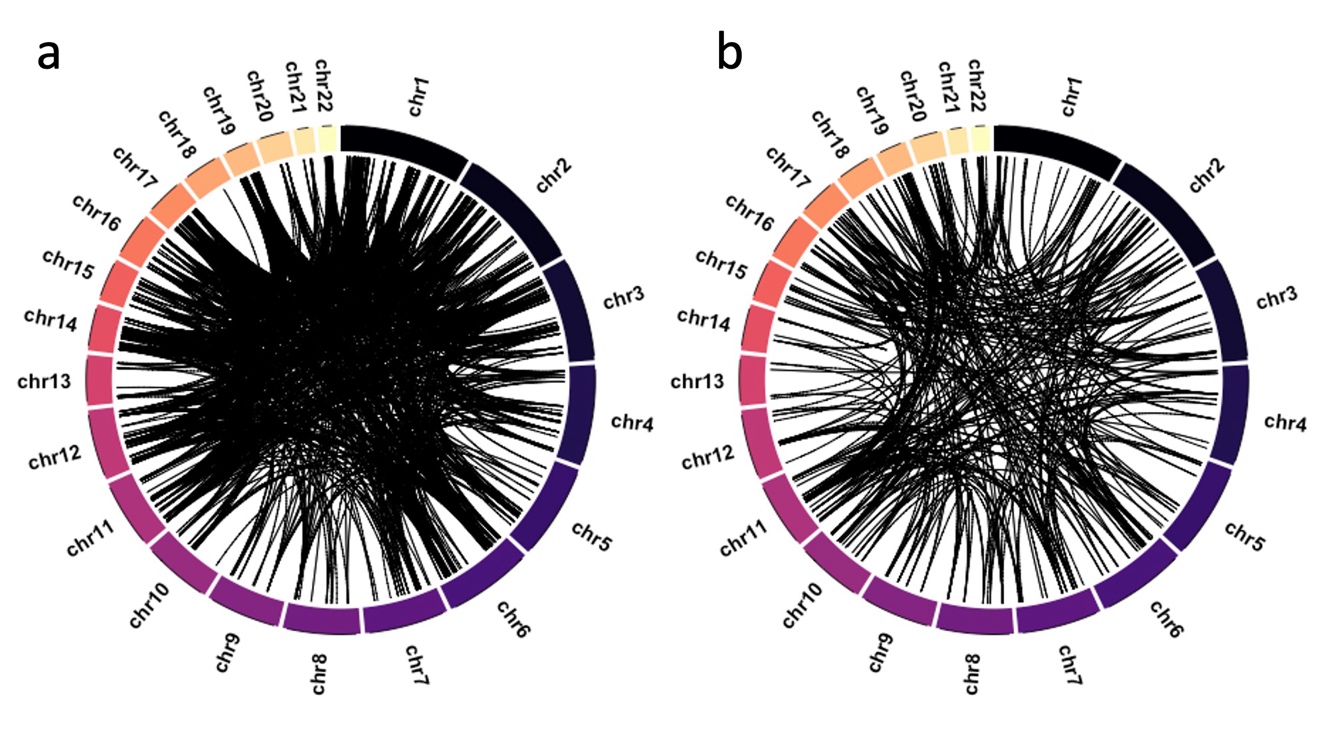

Supplementary Figure 12: Protein–protein interaction (PPI) networks for aSCPs and aVCPs derived from heteroskedasticity analysis.
**(a)** aSCPs-associated protein pairs; **(b)** aVCPs-associated protein pairs.
Only interactions with a confidence score > 0.8 are shown. The aSCPs network exhibits a larger number of protein pairs forming dense local connections, indicating tighter functional integration. In contrast, aVCPs show fewer and more dispersed interactions.

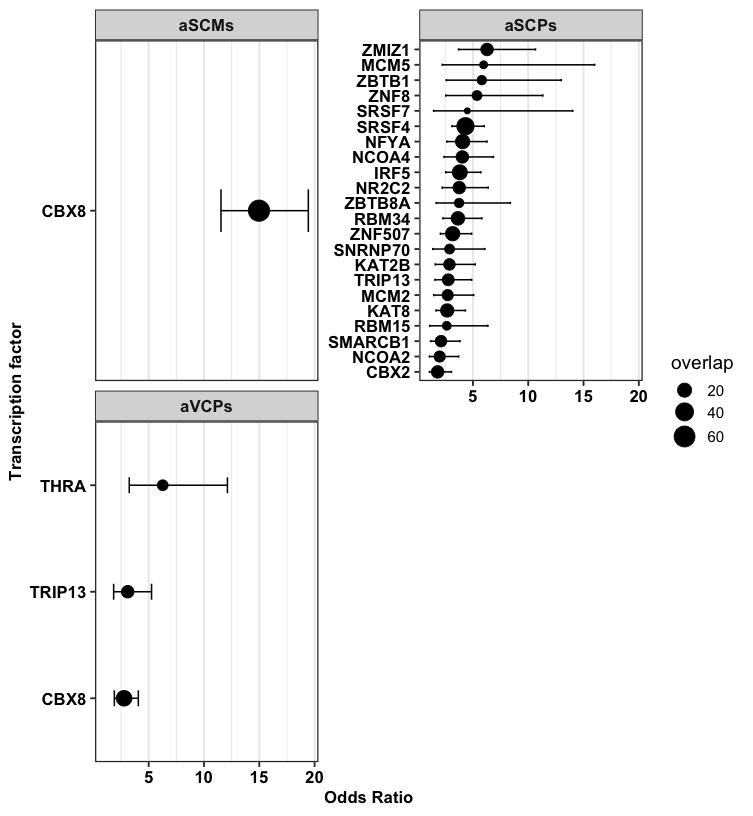

Supplementary Figure 13: Transcription factor binding sites (TFBS) enrichment for age-stable (aSCPs/aSCMs) and age-variable (aVCPs) co-methylation features. Error bars indicate confidence intervals. Only significant hits (p < 0.05) are presented. None of the TFBS reached significance threshold for aVCMs.

Abbreviations: age-stable co-methylation modules (aSCMs), age-variable co-methylation modules (aVCMs), age-stable co-methylated probe pairs (aSCPs), age-variable co-methylated probe pairs (aVCPs).

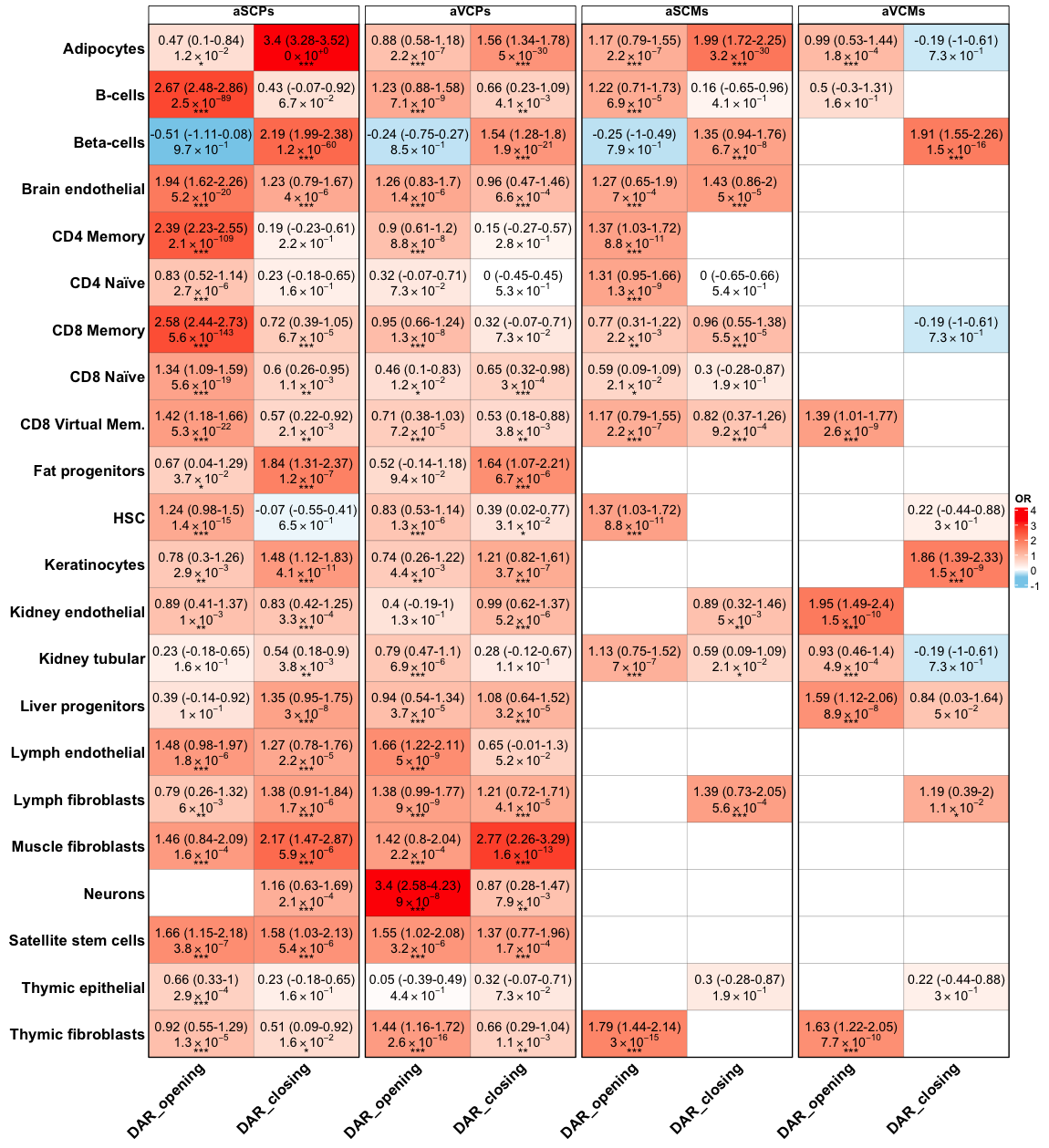

Supplementary Figure 14: Chromatin accessibility enrichment for age-stable (aSCPs/aSCMs) and age-variable (aVCPs/aVCMs) CpG sets.
Differentially accessible regions (DARs) were obtained from Patrick, et al. ^3^, who profiled chromatin opening and closing between young and old mice. The bars are coloured based on log(OR). OR < 0 indicate depletion, while OR > 0, represent over-representation. The asterisks are the significance strength (* = p < 0.05; ** p < 0.01; *** = p < 0.001). The detailed results are given in Table S10.

Abbreviations: age-stable co-methylation modules (aSCMs), age-variable co-methylation modules (aVCMs), age-stable co-methylated probe pairs (aSCPs), age-variable co-methylated probe pairs (aVCPs).

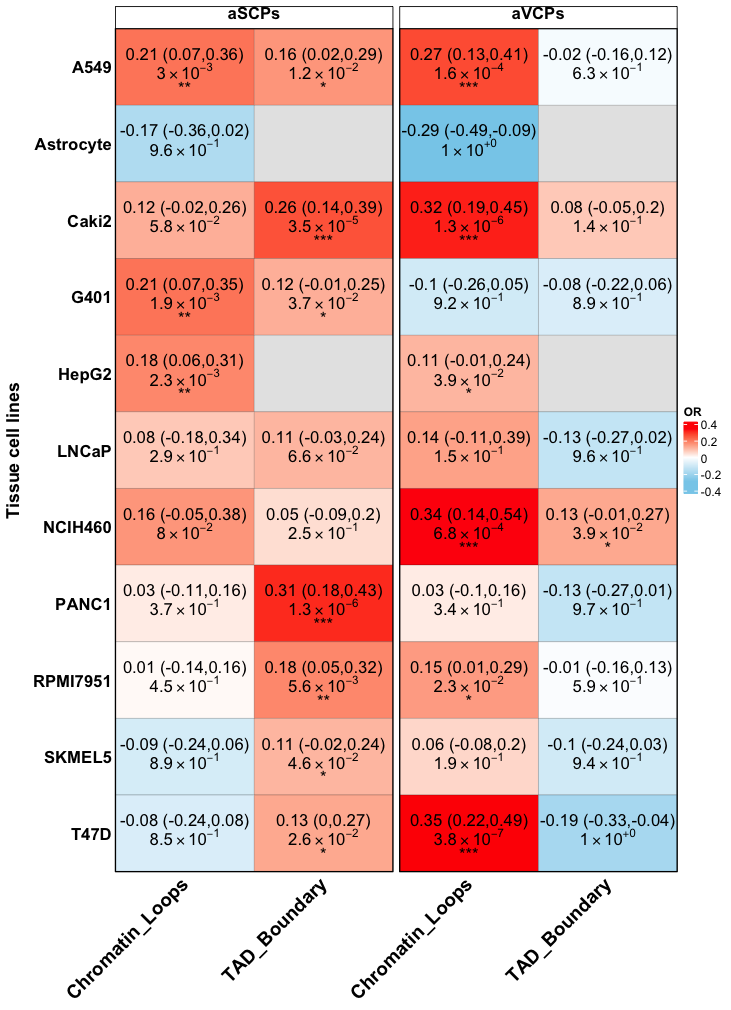

Supplementary Figure 15: Enrichment of age-related pairwise co-methylation sets (aSCPs, aVCPs) within higher-order chromatin architecture features. The bars are coloured based on log(OR). The value in brackets represent the confidence intervals for the OR (in log scale) and the asterisks are the significance strength (* = p < 0.05; ** p < 0.01; *** = p < 0.001). The detailed results are given in Table S11 and S12.

Abbreviations: age-stable co-methylated probe pairs (aSCPs), age-variable co-methylated probe pairs (aVCPs).

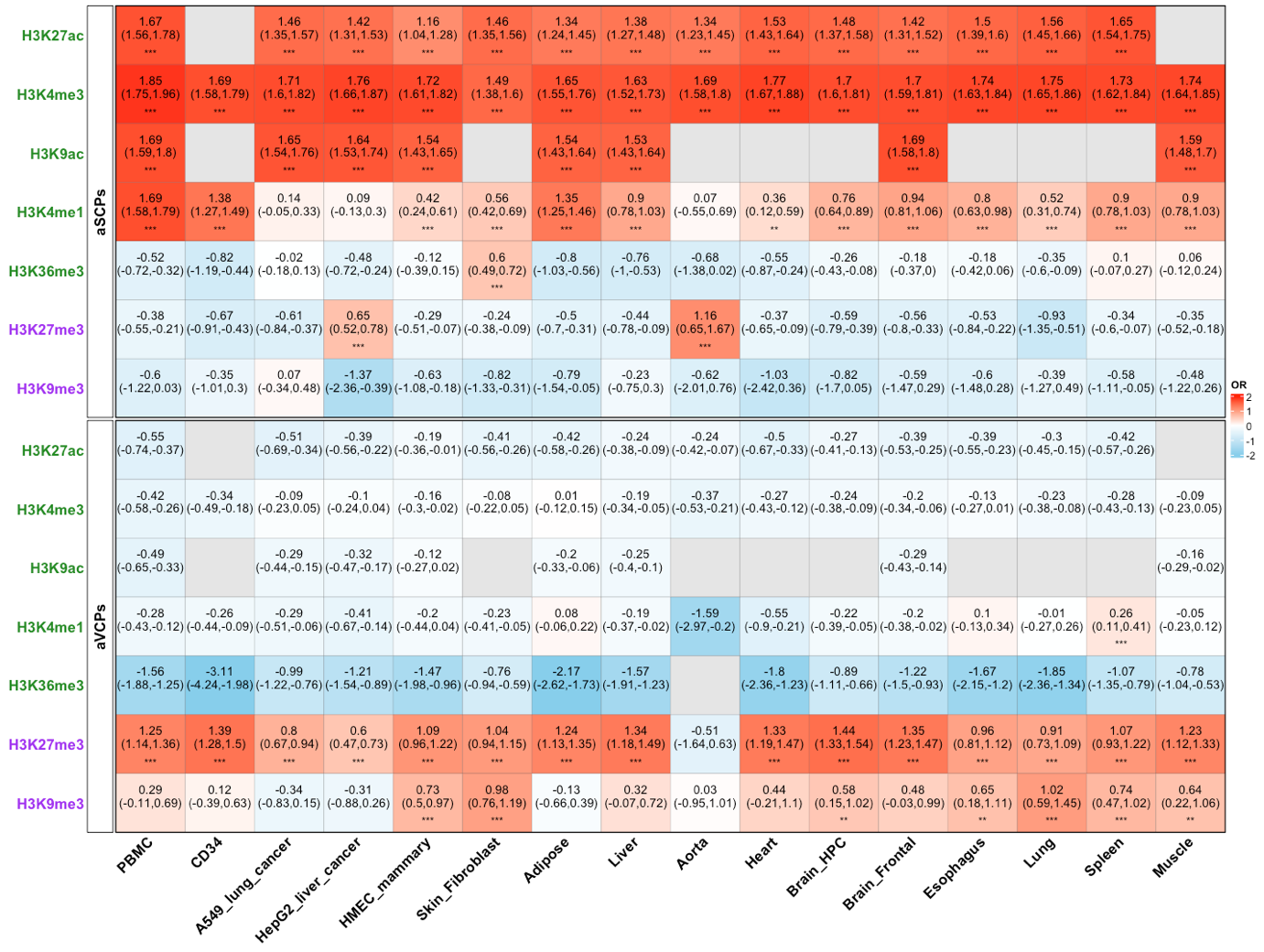

Supplementary Figure 16: Enrichment of aSCPs (Stable) and aVCPs (Changing) for histone modifications across different cell lines.

Histone marks are grouped into active (green) and repressive (purple) categories. CpG sets identified through heteroskedasticity analysis were assessed for enrichment of histone modifications across ENCODE cell lines, categorized as either proliferative or non-proliferative. Overlap enrichment was tested using a one-sided hypergeometric test. The value in brackets represent the confidence intervals for the OR (in log scale), while the asterisks represent significance strength (* = p < 0.05; ** p < 0.01; *** = p < 0.001). OR < 0 and OR > 1, indicate depletion and over-representation of selected age-related sets for specific histone marks, respectively. aSCPs showed strong and consistent enrichment for four active histone marks (H3K27ac, H3K4me3, H3K9ac, H3K36me3) across all tissues. In contrast, aVCPs were consistently enriched for the repressive mark H3K27me3 in most tissues (except for aorta) and displayed modest enrichment for H3K9me3 in several tissues. The detailed results are provided in Table S13.

Abbreviations: age-stable co-methylated probe pairs (aSCPs), age-variable co-methylated probe pairs (aVCPs).

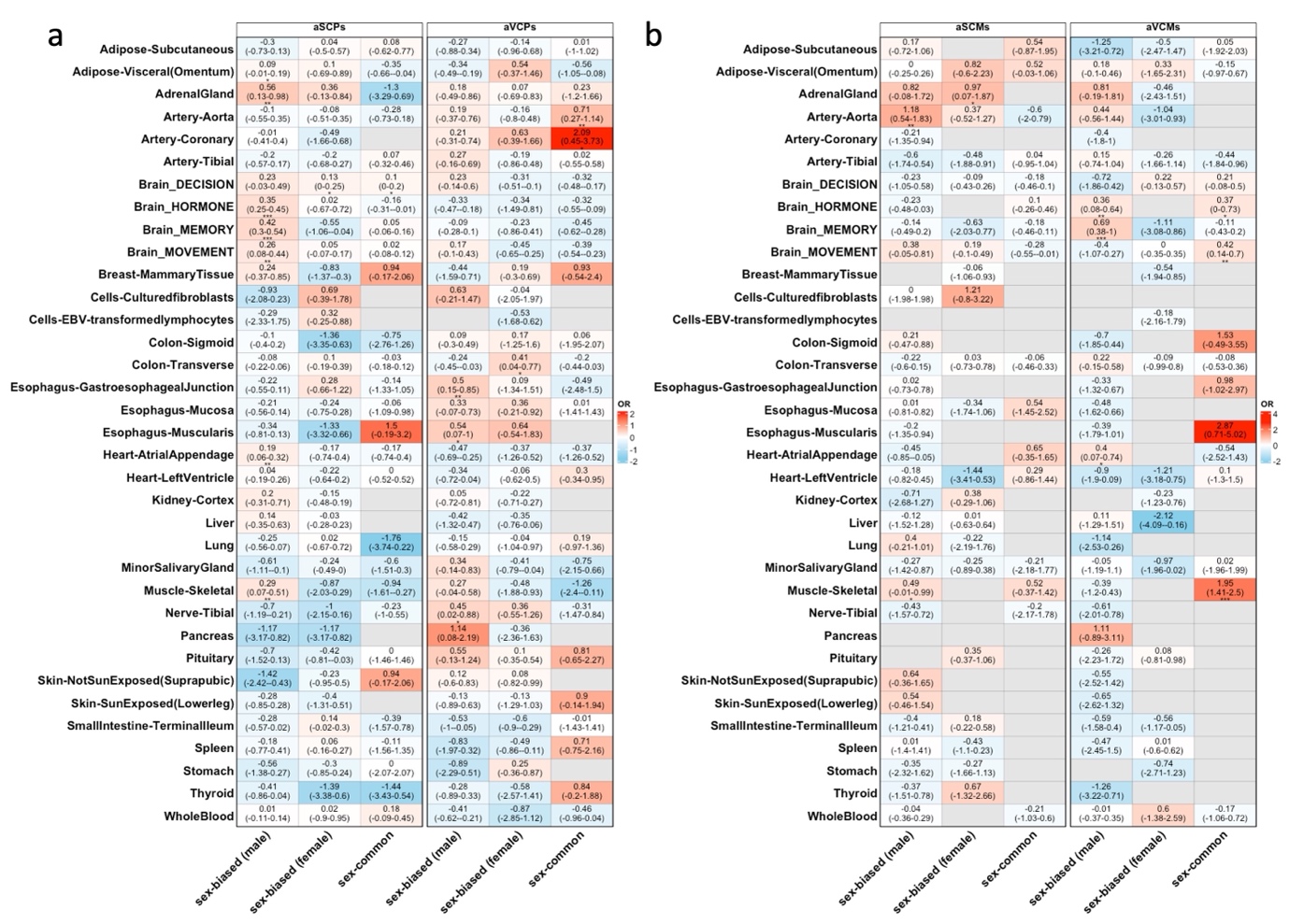

Supplementary Figure 17: Overlap of age-stable and age-variable CpG sets with sex-specific gene expression across tissues.
**(a)** Overlap between aSCPs and aVCPs (derived from heteroskedasticity analysis) and sex-differentially expressed genes across multiple tissues, based on GTEx data from Wang, et al. ^4^. The value in brackets represent the confidence intervals for the OR (in log scale). The grey cells indicate no overlap. Overall, no significant enrichment was observed across most tissues or within either sex, with the exception of few tissues.
**(b)** Same as a but enrichment for aSCMs and aVCMs co-methylation networks. Consistent with (a), no significant overrepresentation was observed across tissues or sexes, with the exception of few tissues.

Abbreviations: age-stable co-methylation modules (aSCMs), age-variable co-methylation modules (aVCMs), age-stable co-methylated probe pairs (aSCPs), age-variable co-methylated probe pairs (aVCPs).

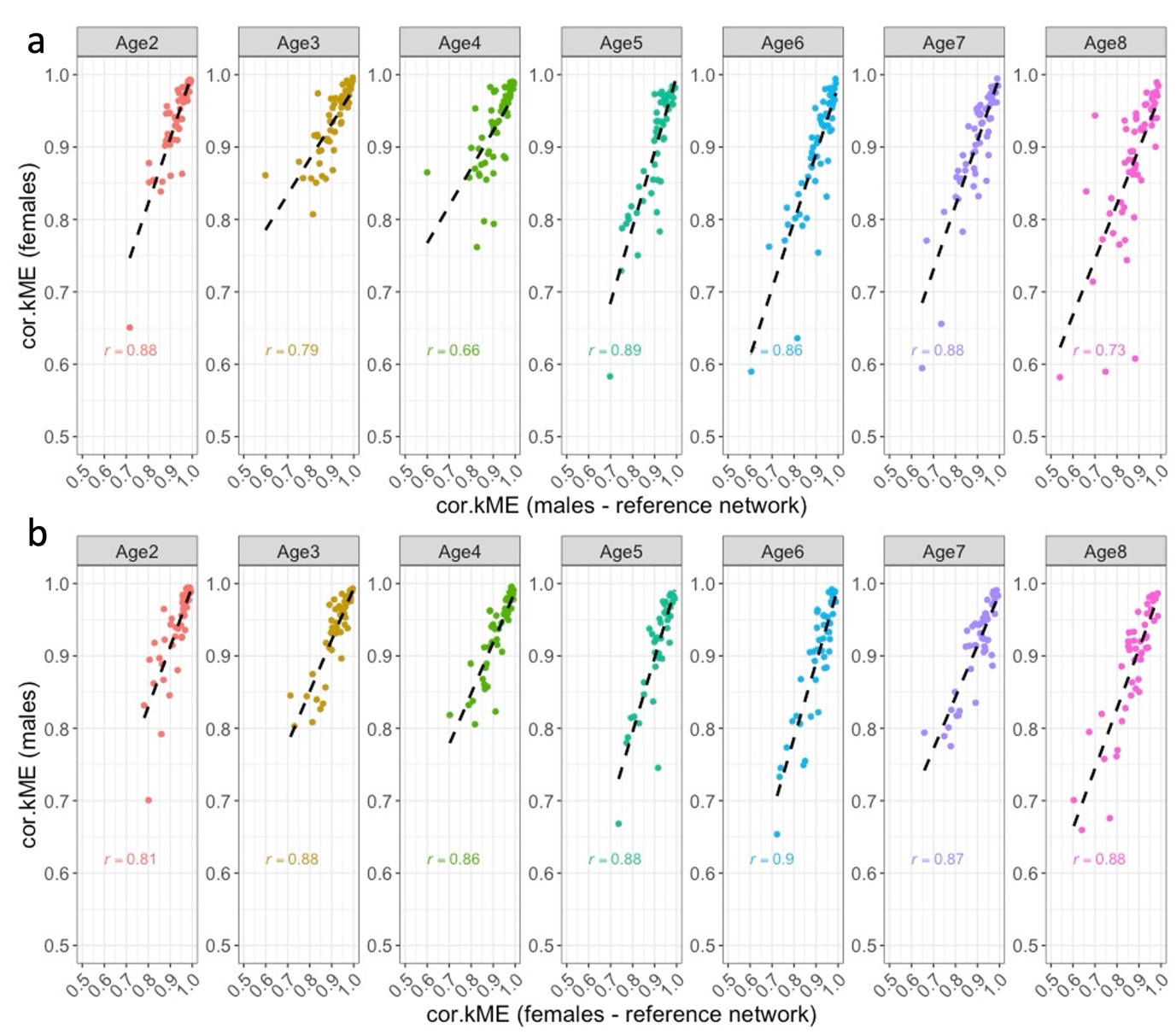

Supplementary Figure 18: Correlation of module preservation statistics

(cor.kME)between males and females across age strata.

**(a)** Male-derived reference networks (N = 84 modules) were constructed using data from the youngest male age group (Age 1: <36 years). Module preservation was then assessed within males and across sex (in females) for each subsequent age group (Age 2 to 8). Each point represents the cor.kME statistic for a single module in males (x-axis) versus females (y-axis).

**(b)** Same as (a) but using female-derived reference networks (N = 79 modules) constructed from the youngest female age group (Age 1: <36 years).

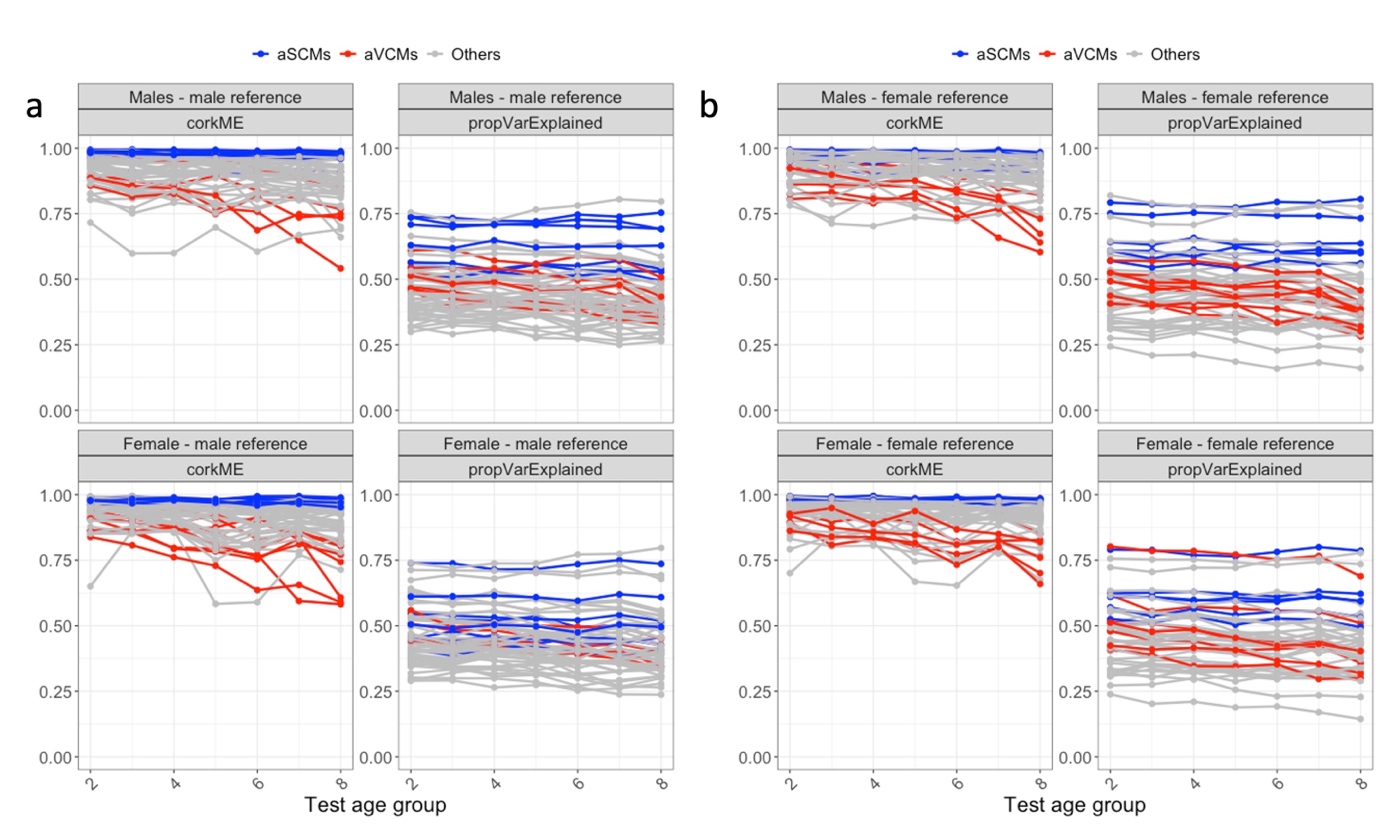

Supplementary Figure 19: Identification of age-stable and age-variable co

methylation modules within sex-stratified networks.

Modules were constructed using the youngest age group (Age group 1, <36 years) for males and females separately, and their preservation values (cor.kME and proportion of variance explained) was tracked across subsequent age groups (Age groups 2–8).
**(a)** Male-derived modules: blue lines indicate age-stable co-methylation modules (aSCMs) with high preservation across age groups; red lines indicate age-variable co-methylation modules (aVCMs) with declining preservation; grey lines represent modules with intermediate or inconsistent preservation patterns.
**(b)** Female-derived modules: classification of modules into aSCMs (blue), aVCMs (red), and other (grey) follows the same criteria as in (a).

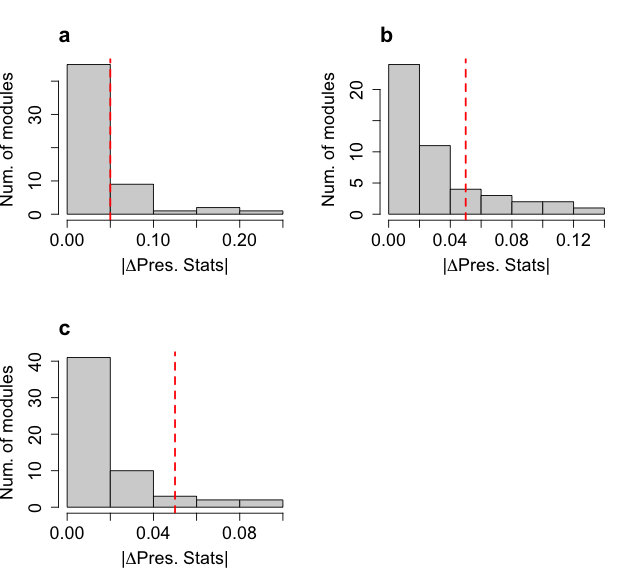

Supplementary Figure 20: Distribution of changes in module preservation

statistics.

**(a)** Change in preservation statistics based on modules derived from the male reference set. For each sex, preservation statistics were compared between extreme age groups (Age Group 2 vs. Age Group 8). The absolute difference in change between males and females was then calculated. The vertical red line (|Δ| > 0.05) corresponding to top 25% of the distribution.

**(b)** Same as (a) but based on modules derived from the female reference set.

**(c)** Same as (a), but the change in preservation statistics was computed between younger and midlife age groups (Age Group 2 vs. Age Group 4), rather than extreme age groups.

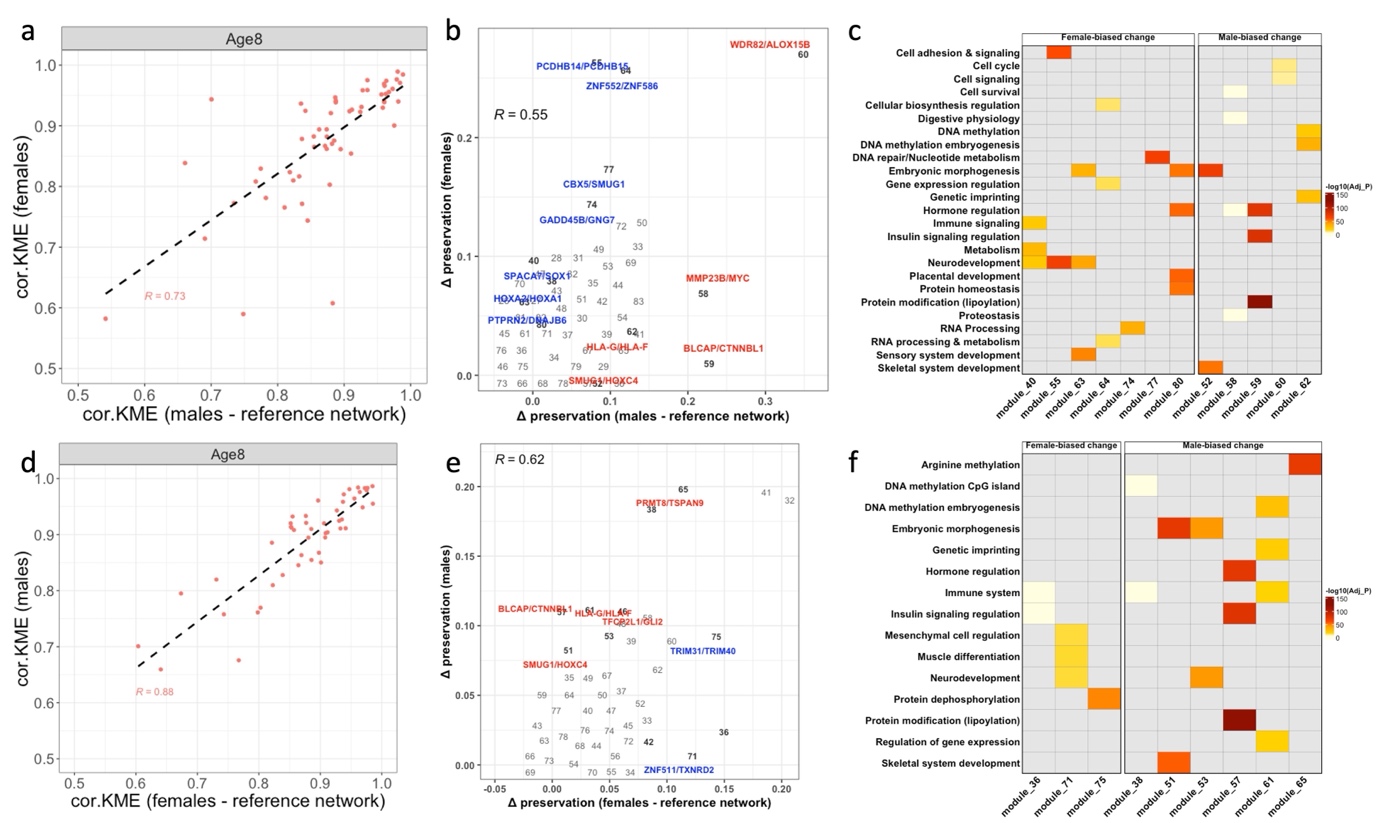

Supplementary Figure 21: Sex-stratified analysis identifies modules with sex-biased age-related co-methylation remodelling.

**(a)** Scatter plot comparing raw network preservation statistics between males and females using male-derived reference networks. Each point represents a module derived from a male-based reference dataset (Age < 36 years).

**(b)** Scatter plot showing absolute changes in module preservation statistics (cor.kME) between younger (36–44 years) and older (>66 years) age groups, computed separately for males (x-axis) and females (y-axis). Modules exhibiting sex-biased preservation changes (>5% difference between sexes) are highlighted and labelled by representative gene clusters.

**(c)** Functional enrichment analysis of sex-biased modules highlighted in (b).
(d, e, f) Equivalent analyses to (a–c), using female-derived reference networks from younger individuals (<36 years).

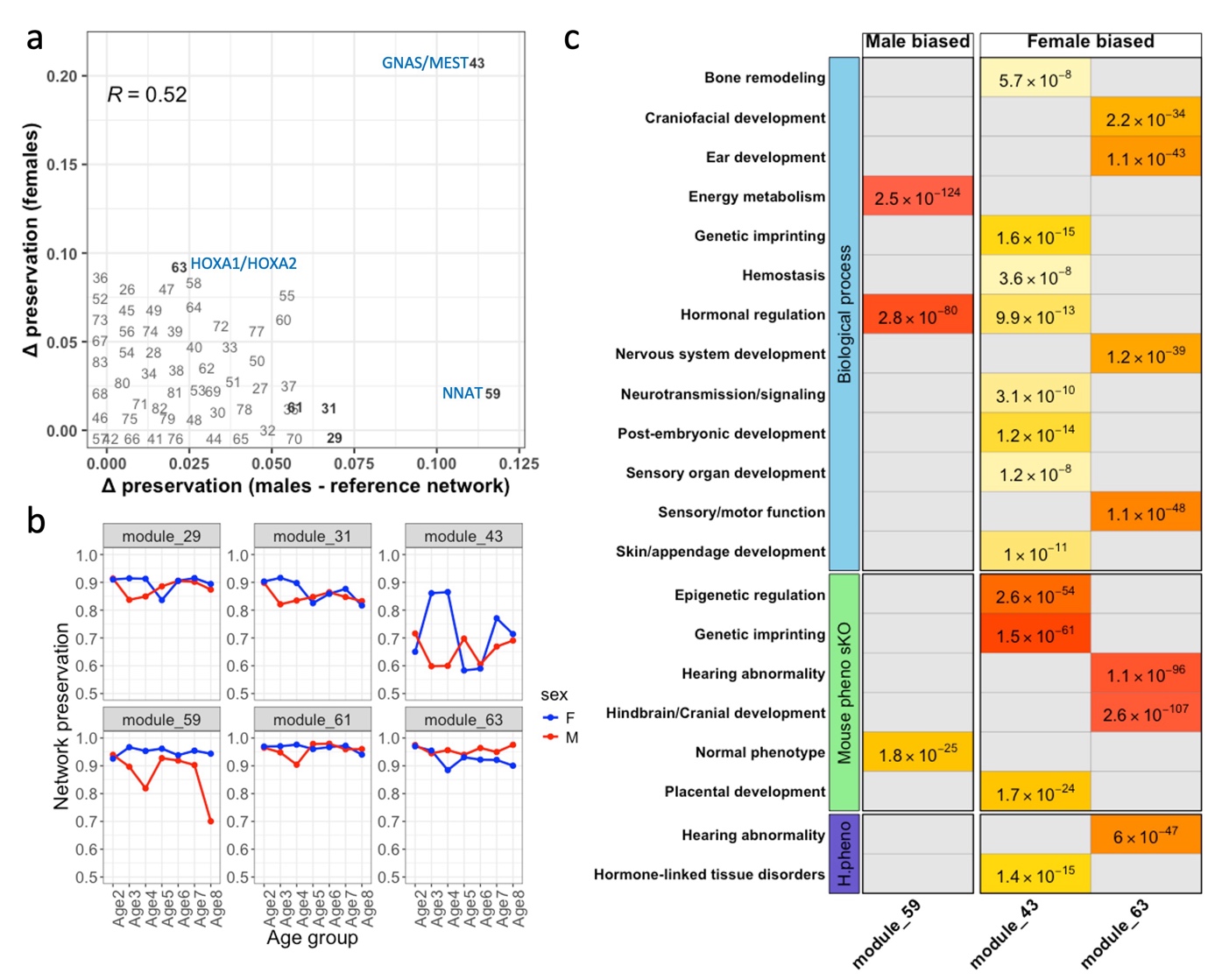

Supplementary Figure 22: Age- and sex-dependent changes in module

preservation and functional enrichment.

**(a)** Scatter plot illustrating the change in module preservation statistics (Δcor.kME)

between age-group 2 (36–44 years) and age-group 4 (50–55 years) in males and

females. Each point represents a module derived from reference data in males <36

years. Modules showing substantial sex-specific change (> 0.05) are highlighted in

bold. Blue labels denote top-ranked genes (p < 0.05) based on rGREAT functional

enrichment. Two modules (43 and 59) exhibit marked divergence in preservation

between sexes.

**(b)** Line plot depicting preservation statistics (cor.kME) across age groups, stratified

by sex, for the sex-biased modules. For module 43, preservation in females peaks at

age-group 4 and then converges with male values. For module 59, preservation in

females remain relatively stable, while in males it gradually declines with age, with

the steepest drop at age-group 8. Overall, sex differences in preservation

trajectories are evident across these modules.

**(c)** Functional enrichment results for the three modules sex biased modules, with statistically significant (p < 0.05) biological processes. Bars are coloured based on -log10(p-values).

**References**

1	Min, J. L. *et al.* Genomic and phenotypic insights from an atlas of genetic effects on DNA methylation. *Nat Genet* **53**, 1311-1321 (2021). <https://doi.org/10.1038/s41588-021-00923-x>

2	Langfelder, P. & Horvath, S. WGCNA: an R package for weighted correlation network analysis. *BMC Bioinformatics* **9**, 559 (2008). <https://doi.org/10.1186/1471-2105-9-559>

3	Patrick, R. *et al.* The activity of early-life gene regulatory elements is hijacked in aging through pervasive AP-1-linked chromatin opening. *Cell Metab* **36**, 1858-1881.e1823 (2024). <https://doi.org/10.1016/j.cmet.2024.06.006>

4	Wang, S., Dong, D., Li, X. & Wang, Z. Pan-tissue Transcriptome Analysis Reveals Sex-dimorphic Human Aging. *bioRxiv*, 2023.2005. 2026.542373 (2023).
